# Tracing genome size dynamics in sharks and rays with inclusive sequence analysis by the Squalomix Consortium

**DOI:** 10.1101/2025.06.08.657570

**Authors:** Shigehiro Kuraku, Yawako W. Kawaguchi, Taiki Niwa, Ryo Misawa, Mitsutaka Kadota, Kazuhiro Saito, Waichiro Godo, Tatsuya Sakamoto, Wataru Takagi, Sachiko Isobe, Kenta Shirasawa, Akane Kawaguchi

**Affiliations:** Molecular Life History Laboratory, National Institute of Genetics, Mishima, Shizuoka, 411-8540, Japan; Department of Genetics, Sokendai (Graduate University for Advanced Studies), Mishima, Shizuoka, Japan; Laboratory for Phyloinformatics, RIKEN Center for Biosystems Dynamics Research, Kobe, Hyogo, 657-0024, Japan; Demersal Fish Resources Division, Fisheries Stock Assessment Center, Fisheries Resources Institute, Japan Fisheries Research and Education Agency, 25-259 Shimomekurakubo, Same, Hachinohe, Aomori, 031-0841, Japan; National Fisheries University, Japan Fisheries Research and Education Agency, 2-7-1 Nagata-honmachi, Shimonoseki, Yamaguchi 759-6595, Japan; Ushimado Marine Institute, Okayama University, Setouchi, Okayama, 701-4303, Japan; Laboratory of Physiology, Atmosphere and Ocean Research Institute, The University of Tokyo, Kashiwa 277-8564, Japan; Kazusa DNA Research Institute, Kisarazu, Chiba 292-0818, Japan

**Keywords:** whole genome sequencing, Hox gene cluster, tandem gene duplication, C-value enigma, ray, cartilaginous fish

## Abstract

Genomes have maintained stable sets of protein-coding genes during evolution, while chromosome organization and genome size vary drastically. Changes in genome size are often attributed to variable amounts of repetitive sequences, including transposable elements. However, it remains poorly understood how such changes were accommodated while maintaining other genomic components. Elasmobranchs, including sharks, rays, and skates, exhibit high among-species variation of genome size and high within-species variation of chromosome length, offering a unique study system to address the question. In this study, we present the first whole genome sequences of the whitebelly skate with remarkably small genome size among elasmobranchs (2.2 Gb), and the red stingray. These chromosome-scale assemblies enabled the assessment of genomic compositions including centromeres and non-coding elements, which revealed notable profiles of tRNA loci and unbiased intragenomic distribution of transposons in elasmobranch genomes. Comparative analyses across these species revealed a shared genomic architecture characterized by correlations of intergenic and intronic sequence lengths with chromosome sizes, with repetitive element accumulation in elongated regions. We analyzed tandemly duplicated genes with high copy number variability. This genome-wide survey revealed the tendency for more frequent tandem gene duplications along with genome size expansion. We document the batoid HoxC cluster in the red stingray genome, which has undergone extensive repetitive element invasion and co-localizes with the HoxB cluster on a sex chromosome. Our study demonstrates an inclusive analysis encompassing both coding and non-coding regions, adaptable to more species in the taxon and a basis for molecular-level understanding on phenotypic diversity of elasmobranchs.

## Introduction

During evolution, genomes have maintained stable sets of protein-coding genes, while chromosome organization—or karyotype—and genome size sometimes underwent drastic changes (Gregory, 2005; Blommaert, 2020). Changes in genome size are often attributed to variable amounts of transposable elements (e.g., Nowoshilow et al., 2018). However, it remains largely unexplored, in a phylogenetic context, how such changes were accommodated in the genome while maintaining complex readout realized by other basic components including protein-coding genes. Cartilaginous fishes, comprising elasmobranchs (sharks and batoids) and chimaeras, exhibit a relatively high genome size variation (Torralba Sáez, et al., 2024) and represent a unique evolutionary lineage within vertebrates, and batoids (rays, skates, torpedoes, and sawfishes; Last et al., 2016) are among the least explored taxa among them at the molecular level (Fig. 1A).

**Figure 1.**
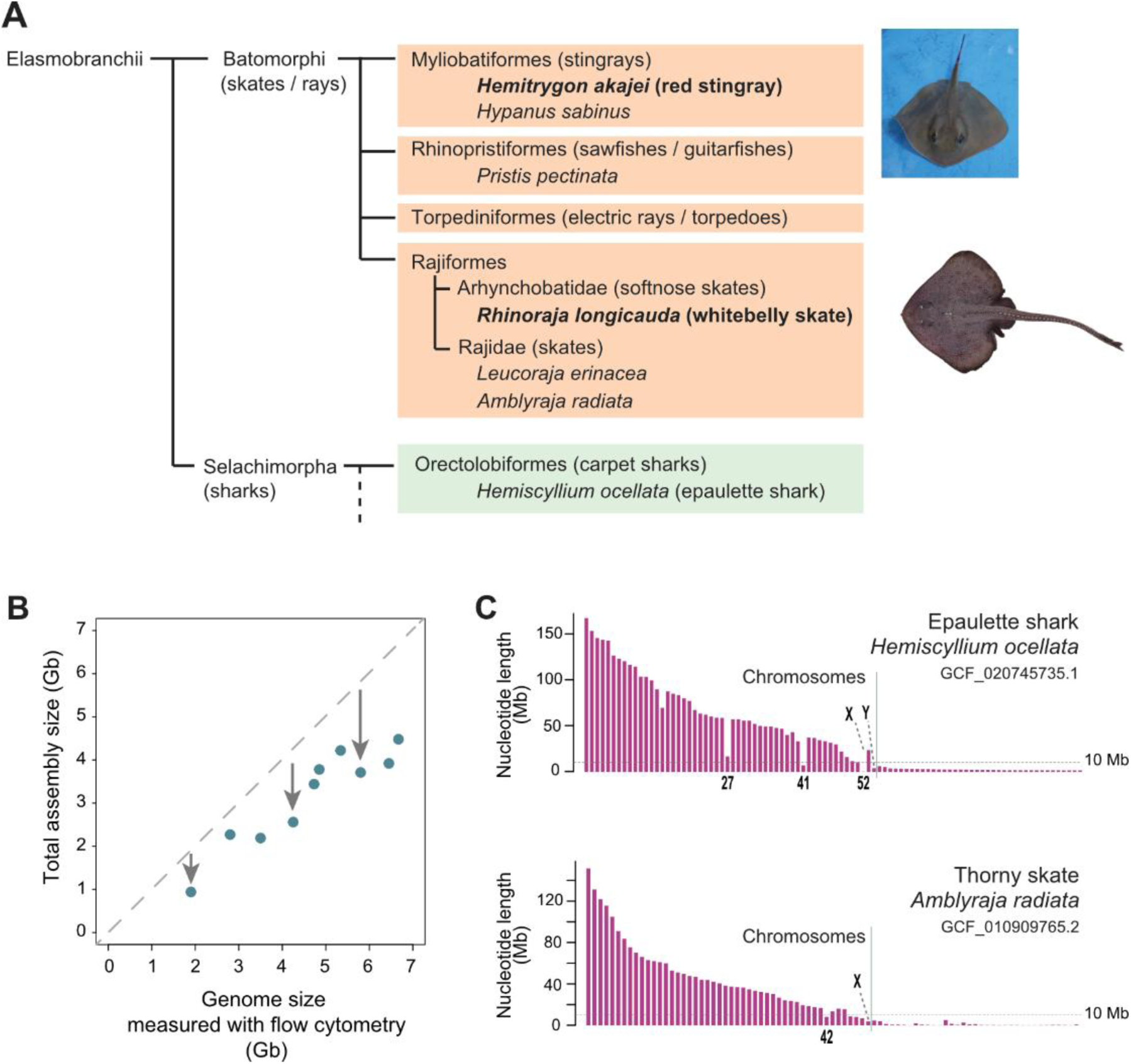
Challenges in elasmobranch genomics in sequence assembly. A, Taxonomy of our study species and their close relatives with available genome assemblies. Photo credits: Shigehiro Kuraku (upper) and Ryo Misawa (lower). B, Systematic underestimate of genome sizes. Original numerical data in this plot are included in Supplemental Table S1. C, Sequence length distribution with ambiguous boundary between ‘chromosomal’ sequences and the other unplaced sequences for two exemplar cases from previously released data (Rhie et al., 2021; Sendell-Price et al., 2023).

Deluge of whole-genome sequencing, propelled by advances in biodiversity genomics (Lewin et al., 2022), has recently been extended to cartilaginous fishes, resulting in the release of dozens of genome assemblies (Wagner et al., 2023; Wu et al., 2024; Yamaguchi et al., 2023). Shark genome analyses have revealed that genome size expansion is driven by intron elongation and the extensive insertion of repetitive elements (Hara et al., 2018; Teramura et al., 2024). Accurate and complete genome assemblies are essential for determining the presence or absence of specific genomic components—e.g., Hox C genes were not identified in genome assemblies for several shark and ray species (reviewed in Kuraku, 2023). Genome-wide studies on the intragenomic distribution of repetitive elements have shown that larger chromosomes tend to accumulate more interspersed repeats, while smaller chromosomes are enriched with short tandem repeats (Yamaguchi et al., 2023). Although genomes in batoids are being released recently, they cover only a narrow subset of its entire diversity (Marlétaz et al., 2023; Rhie et al., 2021; Song et al., 2023; Zhou et al., 2023). It also remains unknown whether this pattern holds in batoids, and whether genome scaling through repetitive element insertions occurs uniformly across different segments of the genome. Additionally, it has not yet been examined whether genome scaling correlates with changes in the abundance of other genomic components, such as protein-coding and non-coding gene copy numbers.

Conventional genome assembly methodologies, designed for broad applications across diverse species, often produce assemblies smaller than the independently measured genome sizes of the same species (Fig. 1B; reviewed in Kuraku, 2021). This size discrepancy is likely attributable to the abundance of repetitive sequences of the genomes; therefore, it will be critical in performing true ‘genome-wide’ analysis focusing on non-coding elements. To encompass non-coding regions, it is essential to generate assemblies with minimal gaps and with sizes comparable to the actual genome sizes. Furthermore, researchers cannot recognize missing parts of the genome unless the genome size is measured independently—this approach is often unfeasible for species with limited sample availability, such as elusive cartilaginous fishes. Another common issue is the difficulty of accurately reconstructing complete chromosomal sequences, resulting sometimes in a considerable number of so-called ‘unplaced’ sequences (Fig. 1C). Such sequences that do not participate in chromosome-scale sequences often include fragments of chromosome segments with sparse signals of chromatin contacts. For instance, the epaulette shark genome, assembled at the chromosome scale, represents a significant resource investment, utilizing ‘trio’ sequencing (Sendell-Price et al., 2023). However, one sequence labeled as a ‘chromosome’ in this assembly was as short as 300 kb (Fig. 1C). Ideally, decisions on which sequences to label as chromosomes should be guided by karyotype reports. In fact, such reports are often unavailable for sharks and rays. This difficulty is critical with these groups of species that tend to have abundant chromosomes in their karyotypes (Uno et al., 2020; reviewed in Stingo and Rocco, 2001). While telomere-to-telomere genome assembly is being achieved in more species, it remains challenging for cartilaginous fishes. This demands a solid technical basis for chromosome-level comparative studies on this taxon.

In this study, we report the first chromosome-scale genome assemblies of two batoid species (Fig. 1A): the red stingray *Hemitrygon akajei*, a myliobatiform ray, and the whitebelly skate *Rhinoraja longicauda*, a deep-sea skate representing the family Arhynchobatidae (softnose skates; Misawa et al., 2020). Our study, conducted under the Squalomix consortium (Nishimura et al., 2022), demonstrates a methodological framework for identifying chromosomal DNA sequences and supplement high-completeness resources for comparative genomic analyses across a broader range of cartilaginous fish taxa. Utilizing these resources, we reveal the patterns of genome evolution in sharks and batoids, highlighting extensive genomic scaling that involves both non-coding and protein-coding regions.

## Results

### Genome sequencing and assembly for unexplored taxa

We report the first whole genome assemblies of two batoid species, the red stingray and the whitebelly skate from the Myliobatiformes and Rajiformes, namely the two most species-rich orders of Batomorphi, respectively. The red stingray genome assembly, consisting of as few as 268 sequences, includes 32 chromosome-scale sequences that amount to 3.68 Gb, resembling its karyotype (2n = 72; Asahida et al., 1987; Fig. 2A) and the nuclear genome content quantified independently (3.74 Gb; Kadota et al., 2023). The whitebelly skate genome assembly includes 44 chromosome-scale sequences that amount to 2.2 Gb (Fig. 2A; Supplemental Figure 1).

**Figure 2.**
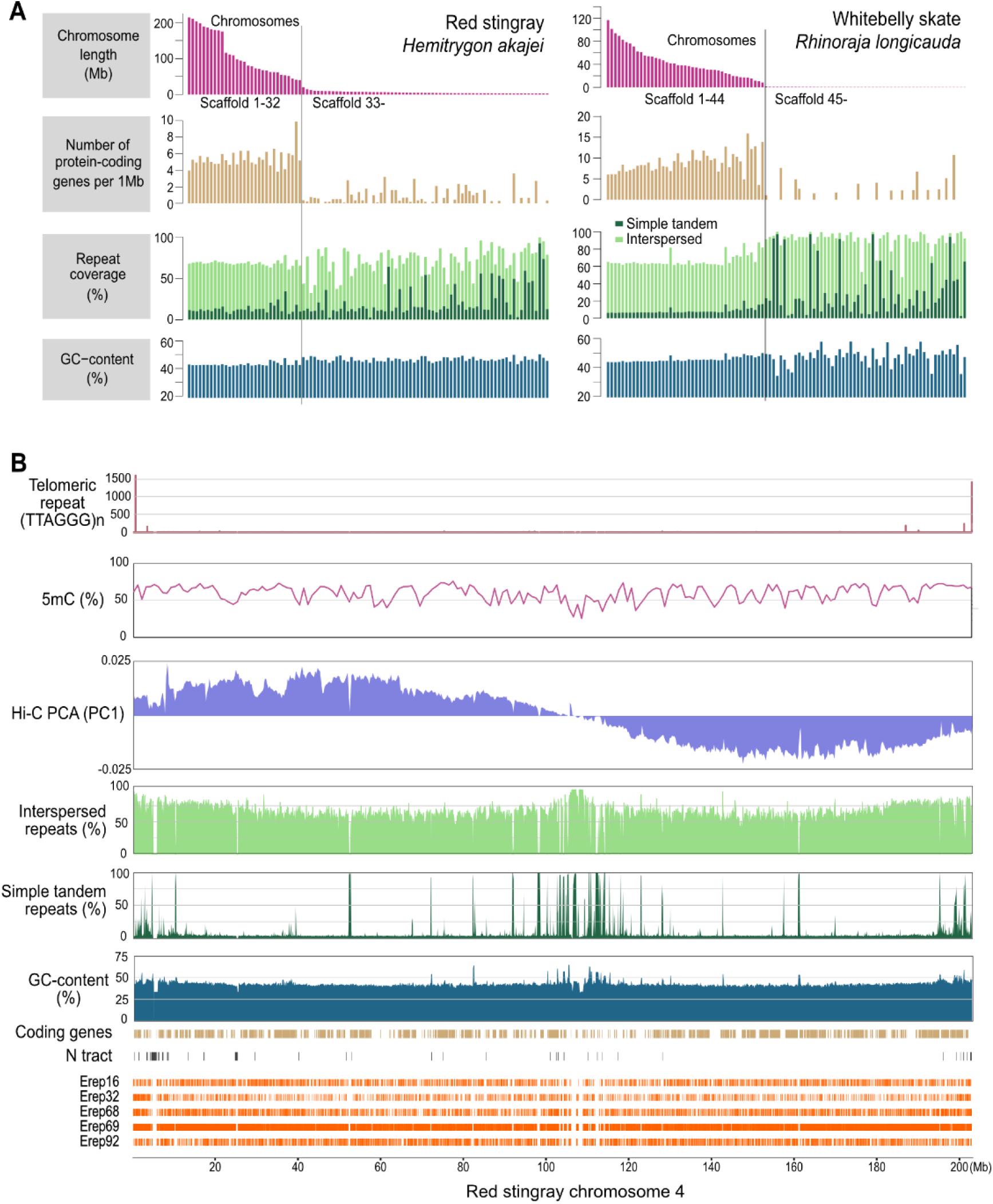
Chromosomal organization in the obtained genome assemblies. A, Chromosome length spectrum instructed by characterization of coding and non-coding genomic landscape. The hundred longest sequences for each species are shown. See Supplemental Fig. S1 for assembly statistics and the comparison with other species. B, Retrieval of the metacentric structure of the red stingray chromosome 4. Proportions of nucleotide bases in interspersed repeats and simple tandem repeats as well as GC bases are shown in 100 kb-long non-overlapping windows, while the frequency of 5-methylcytosine (5mC) in CpG sites is shown in 1 Mb-long non-overlapping windows, based on polymerase kinetics at long-read sequencing (see Methods). Manually curated interspersed repeats abundant in elasmobranch genomes (Erep, elasmobranch repeats; see text) are labeled with their identifiers Erep16 to -92.

Most of these chromosomal sequences have distinguishable features in gene density, repeat abundance and GC-content, apart from their lengths, from non-chromosomal sequences (Fig. 2A). In these assemblies, chromosome-scale sequences occupy large proportions (>90 % with >10 Mb-long sequences), exceeding those of previously reported genome sequences of most shark and ray species (Supplemental Fig. S1). The red stingray chromosomes exhibit a gap in their length distribution between the eleventh largest chromosome (175 Mb) and the twelfth chromosome (116 Mb) (Fig. 2A). The whitebelly skate karyotype resembles that of many shark and batoid species, characterized by abundant chromosomes of highly variable lengths (Fig. 2A).

To characterize the intrachromosomal structure, we focused on the fourth largest chromosomal sequence of the red stingray (Fig. 2B) whose metacentric nature was suggested by an early cytogenetic study (Asahida et al., 1987). Long strings of (TTAGGG)n, a telomeric repeat, were detected uniquely at both ends of this sequence, confirming that it encompassed the whole chromosome. We detected uneven frequencies of interspersed repeats, simple tandem repeats, and GC bases, increasing toward the chromosome ends as well as a central region (positions 105-115 Mb). This central region has the lowest frequency of 5-methylcytosine (5mC) in CpG sites and is marked as a segment across which the frequency of chromatin contacts, captured by Hi-C, is particularly low (Lieberman-Aiden et al., 2009), suggesting it to harbor the centromere (Fig. 2B; see Huang et al., 2023). The retrieval of the putative centromeric structure that remained uncharacterized in earlier studies on cartilaginous fish genomes (Larivière et al., 2024; Rhie et al., 2021) illustrates high completeness of our genome assemblies.

### Retrieving non-coding RNA loci: towards true genome-wide analysis

To shed light on genomic regions largely unexplored in non-osteichthyans, we analyzed loci that produce transfer RNA (tRNA) transcripts. For each of the 20 amino acids, tRNA copies with their corresponding anticodons are transcribed from multiple loci in the genome (Rak et al., 2018; Santos and Del-Bem, 2022). Variable numbers of tRNA loci for the different anticodons comprise a total abundance in the genome that also leads to cross-species differences, previously suggested to be influenced by genome size (Bermudez-Santana et al., 2010). We performed a comprehensive search for tRNA loci in shark and ray genomes by means of primary nucleotide sequence comparison and secondary structure-based validation (see Methods). As a result of our scans, elasmobranchs stood out in the abundance of tRNA loci, with zebrafish being an exception among non-elasmobranch species (Fig. 3A). In particular, shark genomes consistently contained more loci including numerous copies of selenocysteine tRNA that facilitates the use of the 21st amino acid—e.g., 37 copies in the spiny dogfish genome versus only one in all the non-shark species examined (Fig. 3A, B). Our search also revealed the abundance of intron-containing tRNA loci in sharks (Fig. 3A).

**Figure 3.**
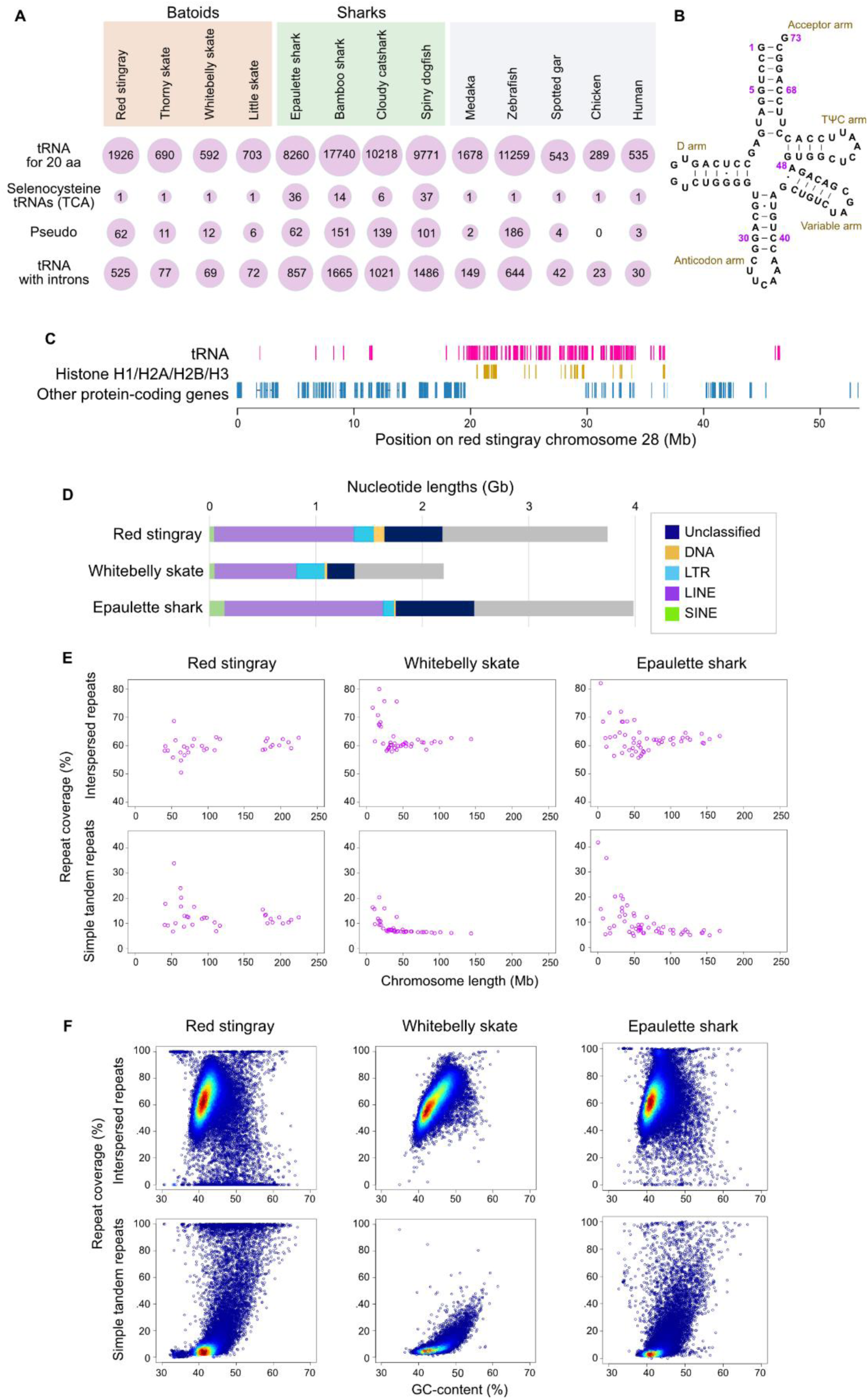
Non-coding landscape in the obtained genome assemblies. A, Quantity of tRNA loci in diverse vertebrates. The size of the circles indicates the per-species abundance in log_10_ scale of tRNA loci in individual categories (see Methods). B, Cloverleaf secondary structure of a spiny dogfish selenocysteine tRNA. This structure, characterized by an exceptionally long variable arm typical of selenocysteine tRNA (Labunskyy et al., 2014), was inferred with R2DT (McCann et al., 2025) using its most abundant nucleotide sequence. C, Locations of tRNA loci on the red stingray chromosome 28 (positions 20-37 Mb) that also contains one of the largest histone gene clusters in this genome (yellow). D, Repeat coverage in the chromosomal sequences of the whole genomes. E, Repeat coverage of individual chromosomes. F, Repeat coverage in different genomic segments in 100 kb non-overlapping windows. See Methods for details on repeat detection.

We investigated the chromosomal locations of the detected tRNA loci, which revealed their biased genomic distribution with clusters on several chromosomes. In the red stingray, chromosome 28 harbors the largest cluster with more than 300 tRNA loci (Fig. 3C). This genomic region also harbors one of the largest clusters of histone H1/H2A/H2B/H3 genes (Fig. 3C), which resembles the cooccurrence of tRNA loci and histone loci on human chromosome 6 and zebrafish chromosome 4 (Reimão-Pinto et al., 2024). The commonality among these species suggests that these two groups of proliferative components (histone genes and tRNA genes) already cooccurred tightly in the genome of their common ancestor more than 450 million years ago.

We recognized that currently available shark and batoid chromosome-scale sequences often lack ribosomal DNA (rDNA) sequences, which are typically challenging genomic regions, as learned from efforts on the human T2T genome (Nurk et al., 2022). In the red stingray genome assembly, we detected regions containing multiple loci of 18S, 5.8S, and 28S rDNA on chromosomes 1, 13, 19, and 20. To facilitate rDNA annotation in other elasmobranch genomes, one of the rDNA sequences detected on red stingray chromosome 20 including its 2kb-long flanking regions on both ends was registered in the sequence database (NCBI Accession ID, LC860076).

### How does repeat invasion influence genome size?

Conventional repeat annotations for the red stingray and whitebelly skate covered 61.2 % and 63.9 % of the whole genome assemblies, respectively (Fig. 3D). These coverages exceeded that of some existing records in elasmobranch genomes including that of the Atlantic stingray, a close relative of the red stingray (approximately 43 % according to Larivière et al., 2024). This difference may be attributable to more complete genome assembly and more exhaustive sequence annotation compatible to this taxon, the central target in the consortium Squalomix (Nishimura et al., 2022). Based on the thorough repeat identification, we compared the profiles of interspersed and simple tandem repeats between individual chromosomes, which revealed the common pattern dependent on chromosome lengths, shared especially between the whitebelly skate and the epaulette shark (Fig. 3E). The red stingray showed an altered pattern, with decreased inter-chromosomal heterogeneity of interspersed repeat coverage (Fig. 3E).

To assess intragenomic heterogeneity irrespective of karyotypic differences, we focused on GC-content and repetitiveness that are associated with transcriptomic and epigenomic readout of the genome (Hara and Kuraku, 2023). Our genome-wide profiling of GC bases and repetitive elements revealed a remarkable tendency for highly repetitive regions to have higher GC-content (Fig. 3F). The whitebelly skate, with a relatively small genome among elasmobranchs, has much fewer genomic regions occupied predominantly by simple tandem repeats (e.g., 50-100 %), than the red stingray and epaulette shark (Fig. 3F). This also explains the abundance of regions with low interspersed repeat coverage (e.g., 0-30 %) in the red stingray and epaulette shark (Fig. 3F), assuming that those regions are occupied by simple tandem repeats that are also abundant in these species.

To further characterize the repetitive landscape, we focused on repetitive element models with high abundance in the red stingray genome. From the 1,506 elements in the *de novo* repeat library, we selected 10 elements with two criteria: the length of >2 kb and the matching regions of >50,000 in the red stingray genome. As demanded generally for high-accuracy repeat annotation (Goubert et al.,2022; Platt II et al., 2016), we performed manual sequence curation of these elements to derive five confident models representing >1,000 high-similarity copies detected in this genome (Supplemental Table S2). They scatter throughout the genome except some regions with atypical sequence characteristics including centromeric regions, as shown for chromosome 4 (Fig. 2B). Their unbiased distribution suggests no overt regional bias as a consequence of insertions and maintenance of these repetitive elements.

### Which contributed more to genome expansion, intronic or intergenic regions?

Previously, intron elongation was shown to contribute to genome size enlargement (Hara et al., 2018; Fig. 4A). The comparison involving the species analyzed in this study suggested that overall change of genome size is subdivided into chromosome-level change of sequence length (Fig. 4B). Still, it has not been examined which portion of the genome, e.g., introns or gene intervals (‘intergenic regions’), has contributed more to changes in genome size, and whether genomic spacing also acts in intergenic regions. First, we compared the total lengths of introns and intergenic regions among species, which revealed the proportional changes of both introns and intergenic regions along with the change in genome size (Fig. 4C). Second, we compared the length distributions of introns and intergenic regions between individual chromosomes in each elasmobranch genome. Longer introns were harbored by genes located on larger chromosomes (*p* < 0.01), although this trend was weakened in the red stingray (*p* = 0.0201; Fig. 4D). A similar trend was observed in the intergenic regions (Fig. 4E). These associations are consistent with the correlation between the lengths of introns and intergenic regions, at the chromosomal level (Fig. 4F).

**Figure 4.**
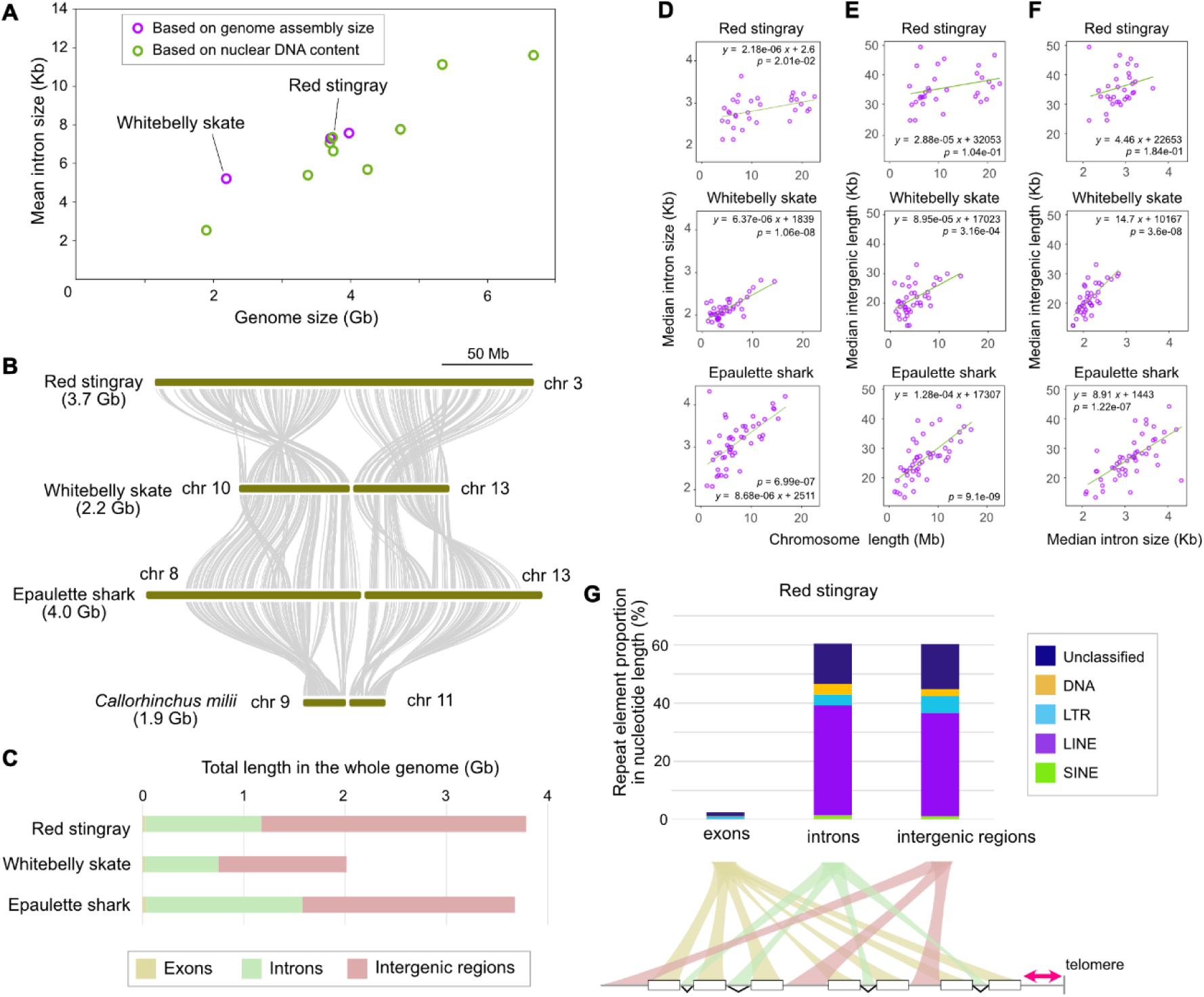
Regional and compositional dissection of genomic spacing. A, Relationship between genome size and intron size. Genome sizes are based on the existing literature (Gregory, 2002; Kadota et al., 2023). Original numerical data in this 2D plot are found in Supplemental Table S3. B, Cross-species size change of homologous chromosomes. Curves between chromosomes show one-to-one orthology of protein-coding genes. C, Breakdown of whole genomes in nucleotide length into exonic (beige), intronic (lightsage), and intergenic (plum) regions. D, Relationship between chromosome length and intron length. Outliers were removed using the interquartile range method. Green lines represent the simple regression. E, Relationship between chromosome length and intergenic length. F, Relationship between intron length and intergenic length. Intronic and intergenic lengths are shown as medians for individual chromosomes. G, Comparison of the landscape of interspersed repeats between exonic, intronic, and intergenic regions. Note that non-genic regions on chromosome ends (shown in red) are not employed particularly for this repetitive element profiling.

In elasmobranch genomes, the extent of repeat accumulation is a strong determinant of their genome size (Fig. 3D; Hara et al., 2018). To further examine which portion of the genome preferentially underwent repeat accumulation, we analyzed the red stingray genome (Fig. 4G), which showed no overt difference between intronic and intergenic regions, whereas exonic regions exhibited much lower coverage of repetitive elements as expected.

### Profiling tandem gene duplications: reflection of genome size?

Genome enlargement has repeatedly been shown to be associated with expansion of non-coding elements such as transposable elements (Nowoshilow et al., 2018; Schartl et al., 2024), but its influence on protein-coding regions has not been explicitly documented (see Kapusta et al., 2017). To address this, we performed a genome-wide profiling of tandem gene duplications. Importantly, investigation of relative gene locations can often be confused by false gene prediction. To overcome such problems and examine tandem gene arrays in a comprehensive manner, we formulated a metric *M_g_* that reflects the abundance of homologous genes surrounding every locus (see Methods). This metric is not largely influenced by intergenic sequence length and false positive gene prediction (causing noise in the detection of gene clusters). With this method, we comprehensively identified regions containing tandem gene copies in the genomes sequenced in this study (Fig. 5A; Supplemental Fig. S3). The results included the family of genes encoding succinate receptors (SUCNR1) with two lineage-specific duplicates in elasmobranchs (Fig. 5B). This method also identified tandem gene clusters previously reported in elasmobranch genomes including protocadherin (Pcdh; Hara et al., 2018), oxytocin/vasopressin (Hara et al., 2018), VLDLR (Ohishi et al., 2023), Lefty (reviewed in Kuraku, 2024), γ-crystallin (Mayeur et al., 2024), and Wnt1-6b-10b (reviewed in Kuraku, 2021), suggesting the reliability of this metric.

**Figure 5.**
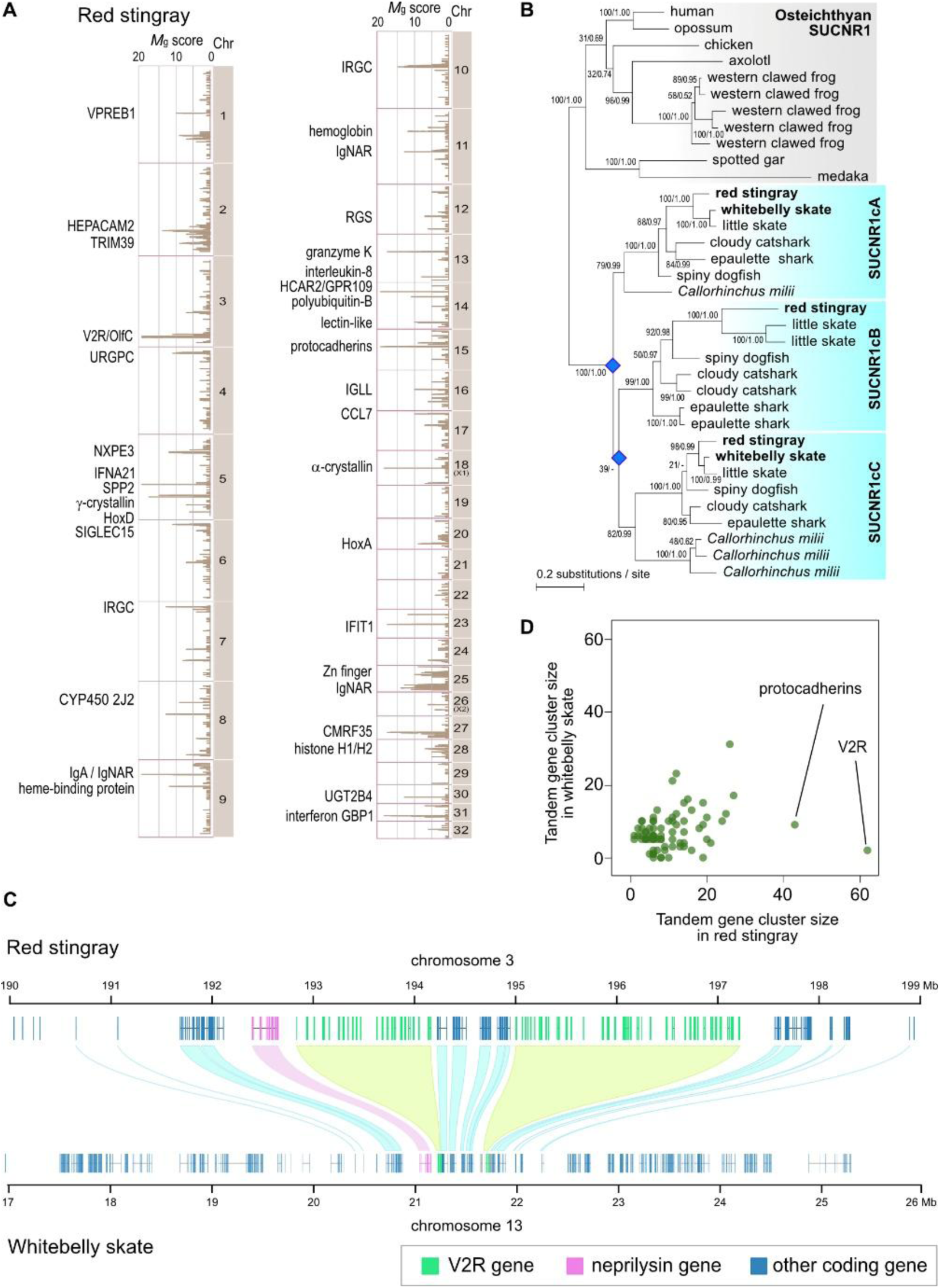
Tandem gene clusters in batoid genomes. A, Landscape of tandem duplicates in the red stingray genome. The *M_g_* scores for individual predicted genes are shown. Names of genes or gene groups are displayed when explicit non-elasmobranch homologs are identified. See Supplemental Table S4 for abbreviations of gene names. B, Molecular phylogeny of succinate receptor gene family (SUCNR1 or GPR91). Blue rhombi show gene duplications in the chondrichthyan lineage. See Supplemental Table S6A for accession details for the sequences used. C, Scaling in the tandem V2R gene clusters. Green areas show cross-species correspondence of the V2R gene clusters, while light blue ribbons indicate one-to-one orthology in other gene families. Pink boxes indicate the exons for the neprilysin gene (consisting of 22 exons) previously shown as a consistent marker adjacent to the orthologous V2R cluster in diverse vertebrate genomes (see text). D, Comparison of the numbers of component genes in the detected tandem gene clusters between the two batoid genomes. Only tandem gene clusters with no less than five component genes in either species are taken into account (81 cases).

The comparison of tandem gene arrays between two batoid species highlighted gene families with dramatic size discrepancy (shown in red in Supplemental Fig. S3), including the HoxC cluster (see below) and vomeronasal type 2 receptors (V2R or OlfC; Silva and Antunes, 2017). In the red stingray genome, we identified the V2R gene cluster with the largest number of components (65 genes) among cartilaginous fishes (Fig. 5C; Syed et al., 2023). In its orthologous chromosomal region on the whitebelly skate, we identified only two V2R homologs (Fig. 5C), including which its entire genome contained as few as seven genes. As shown previously for other vertebrates (Hashiguchi and Nishida, 2009; Silva and Antunes, 2017; Zhang et al., 2022), the V2R-containing region is marked by the adjacent neprilysin gene in both batoid species (Fig. 5C), confirming the orthology of these gene groups across different vertebrate lineages. The whole V2R cluster occupies more than 3 Mb in the red stingray genome, while the whitebelly skate counterpart 0.3 Mb. The shrunken whitebelly skate V2R-containing regions are not interrupted by any undetermined gaps in the sequences, refuting the possibility of incomplete sequencing or assembly. Similarly, the genomic regions flanking the V2R gene cluster are elongated in the red stingray genome (Fig. 5C).

Overall, tandem gene clusters were stably positioned relative to the conserved synteny of neighboring one-to-one orthologs between the two batoid species (Supplemental Fig. S3). The red stingray genome contained 24 regions with loci exhibiting a *M*_g_ greater than 10, compared to only 11 such regions in the whitebelly skate genome. Across the genome, the red stingray harbored significantly more component genes in tandem duplicate clusters than the whitebelly skate (Wilcoxon signed-rank test, *p* = 0.0008), with particularly pronounced expansions in the Pcdh and V2R gene families (Fig. 5D).

### Identifying batoid Hox C: what makes it highly elusive?

Most jawed vertebrates analyzed to date possess four Hox gene clusters—HoxA through HoxD—each typically confined within a genomic region of approximately 100 kb (Dechamps and Duboule, 2017). Still, no batoid species has ever been documented to possess Hox C genes, firmly with phylogenetic and transcriptional evidence. The obtained red stingray genome assembly included a chromosome-scale scaffold sequence (chromosome 18) containing a cluster of HoxC genes, in addition to HoxA, -B, and -D clusters (Fig. 6A). This is the first identification of the batoid HoxC cluster containing the divergent *HoxC3* ortholog that remained unidentified in elasmobranch species (Supplemental Fig. S4B). The deduced *HoxC3* amino acid sequence underwent some substitutions unique to elasmobranchs even in the homeodomain (Fig. 6B). The high sequence divergence of myliobatiform *HoxC3* was also indicated with the elongated branches in molecular phylogeny (Fig. 6C; Supplemental Fig. S4A). The HoxC clusters of the red stingray and other elasmobranch species have fewer Hox genes than in many non-elasmobranch species (Supplemental Fig. S4B; reviewed in Kuraku, 2023). In contrast, HoxA and -D clusters have maintained the gene compositions during shark and ray evolution, exhibiting no gene gain or loss (Supplemental Fig. S5).

**Figure 6.**
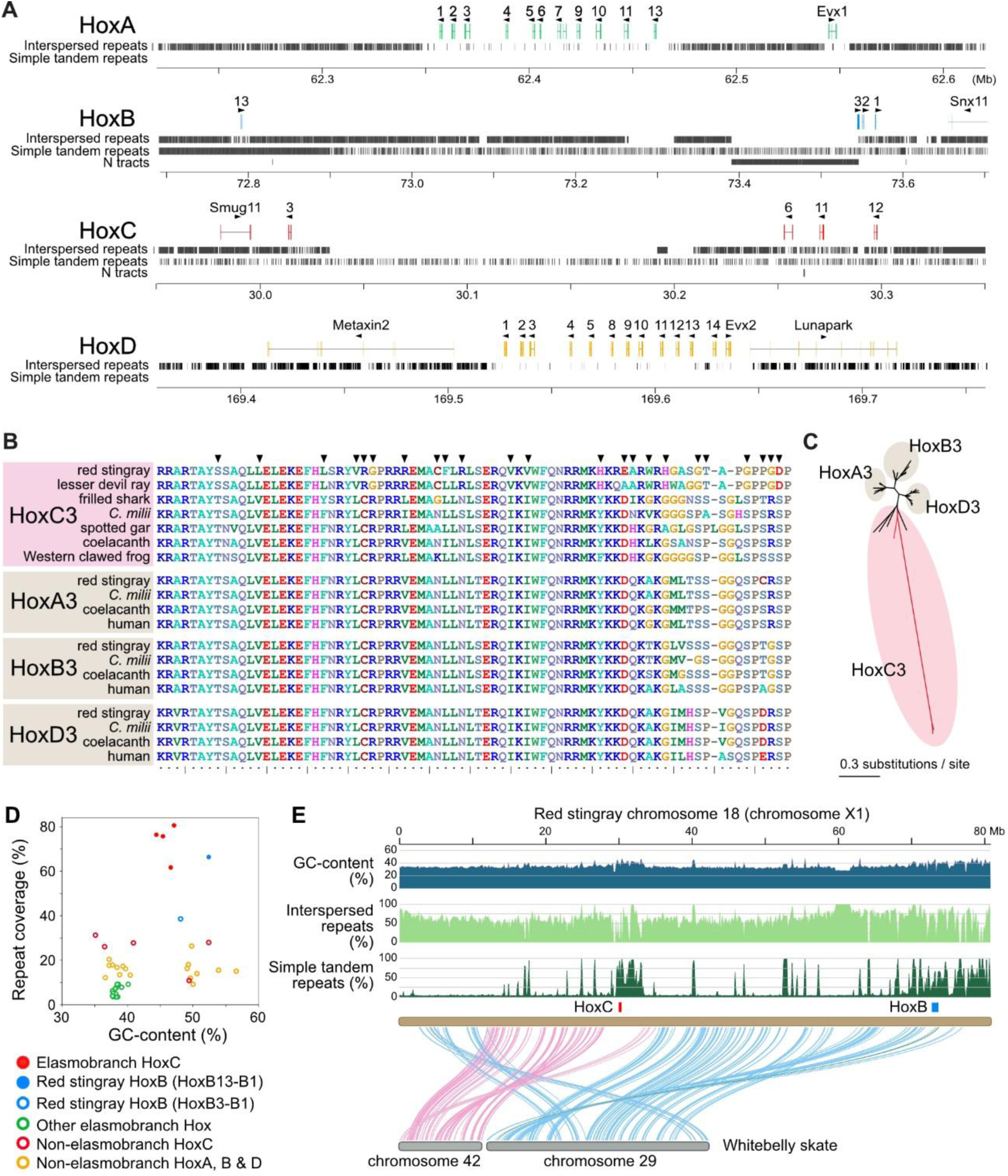
Elasmobranch Hox gene identification. A, Genomic structure of the red stingray Hox clusters and their neighboring regions. The exons of the Hox genes are shown in colored boxes. The genomic sequences containing HoxA and -D clusters (on chromosome 20 and 5, respectively) shown here are interrupted by no gaps in the genome assembly. HoxB and -C clusters are both located on chromosome 18. B, Amino acid sequence alignment of Hox3 genes. Arrowheads indicate the amino acid sites at which HoxC gene products have unique residues. C, Molecular phylogeny of jawed vertebrate Hox3 genes. Red branches show the lineages leading to elasmobranch *HoxC3*. See Supplemental Fig. S4A, for details. D, GC-content and repeat coverage of Hox clusters in different groups of species. Both interspersed and simple tandem repeats are regarded collectively as repeats in this analysis. The nucleotide sequences between the start codon of Hox13 (Hox10 for the B cluster) and stop codon of Hox1 were compared. The numerical data used to form this plot are provided in Supplemental Fig. S6. E, Chromosome-wide structure of red stingray chromosome 18 that harbors both HoxB and -C clusters. Curves connecting with two homologous whitebelly skate chromosomes show one-to-one orthology of protein-coding genes.

We compared the sequence properties of HoxC-containing genomic regions including non-coding sequence segments, which shows outstanding abundance of repetitive elements and higher GC-content, compared with the other Hox clusters (Fig. 6D)—the HoxC cluster seems to have a high level of repetitiveness comparable to the non-Hox genomic regions (Fig. 3B) and to have escaped from high structural constraint observed typically in gnathostome Hox clusters. Our embryonic transcriptome data of the red stingray showed the transcription of all the identified HoxC genes, namely *HoxC3*, *-C6*, *-C9*, *-C11*, and *-C12* (Supplemental Fig. S4C).

Chromosome 18 of the red stingray harbors a HoxB gene cluster as well as the abovementioned HoxC cluster, as a result of a fusion of two chromosomes (Fig. 6E). In contrast, in all the analyzed species except myliobatiforms, the HoxB and HoxC clusters are consistently located on separate chromosomes (Supplemental Fig. S4B). The red stingray genome sequence containing the HoxB cluster harbored four Hox genes— *HoxB1*, *-B2*, *-B3*, and *-B13*—arranged in consistent transcriptional orientations and separated by a stretch of over 700 kb devoid of predicted protein-coding genes (Fig. 6A). This cluster is flanked by orthologs of genes such as *Snx11* and *Tax1bp1*, which similarly flank HoxB clusters in other elasmobranch species, including the whitebelly skate. Notably, the red stingray HoxB cluster, as well as HoxC, exhibits high GC-content and repeat coverage, features that persist even when accounting for the highly repetitive region between *HoxB13* and its neighboring Hox gene (Fig. 6D). The high repeat coverage may explain the absence of certain HoxB genes in the genome assembly, although they were identified in transcriptome data (Supplemental Fig. S4C). These findings align with observations for the genome assembly of the Atlantic stingray (*Hypanus sabinus*, NCBI GCF_030144855.1; Larivière et al., 2024; Lee et al., 2025) which was previously classified in the same genus as the red stingray. In the Atlantic stingray assembly, sequences containing HoxB genes are assigned to chromosome X1 (NW_026778977.1 ‘SUPER_X1_unloc_8’ which is 819 kb-long and NW_026778963.1 ‘SUPER_X1_unloc_15’ which is 541 kb-long), although they remain ‘unplaced’ and unassembled into a chromosomal sequence. Taken together, these observations support the co-localization of the HoxB and HoxC clusters on chromosome X1, uniquely in the myliobatiform species of batoids (Supplemental Fig. S4B).

Exploiting the utility of Hi-C data obtained for genome scaffolding, we identified two topologically associating domains (TADs) in the HoxD-bearing region in the red stingray, with the HoxD cluster positioned at their boundary (Supplemental Fig. S8A). A similar arrangement was observed for the HoxA cluster, which is also located at the interface between two TADs (Supplemental Fig. S8A). We also attempted to analyze chromatin contacts in genomic regions containing the HoxB and -C clusters, but our Hi-C data could not be mapped properly due to the high abundance of repetitive sequences (Fig. 6A; Supplemental Fig. S7). This resulted in low coverage in these regions, which interfered the extraction of interpretable chromatin contacts.

## Discussion

We obtained chromosome-scale genome assemblies of two batoid species and conducted a comparative analysis involving them. Of these, the red stingray is one of the first species to have genome-wide DNA sequences in Myliobatiformes, one of the four batoid orders. The other species whitebelly skate in the order Rajiformes provided one of the smallest genome assemblies among Elasmobranchii as of April 2025 (Supplemental Fig. S1). We used these novel assemblies as main materials for establishing an analysis standard for elasmobranch genomes spanning telomeres, centromeres, rDNAs, tRNAs, and repetitive elements as well as protein-coding genes. Recent studies in various taxa illustrated the contribution of massive interspersed repetitive elements to genome size expansion (Brown et al., 2025; Nowoshilow et al., 2018; Schartl et al., 2024), a pattern also observed in sharks (Hara et al., 2018). However, how genome size changes occur and how different genomic components are affected have remained unclear. Our dataset, including a species pair with contrasting genome sizes, provided a valuable basis for addressing these questions.

Our cross-species comparisons revealed that larger genomes tend to harbor more tandem copies of protein-coding genes and putative tRNA loci, as well as longer intronic and intergenic sequences (Figs. 3, 4, and 5). Importantly, the abundance of the detected tRNA loci persisted even with a stringent condition of tRNA detection (see Methods). tRNAs are known to derive PIWI-interacting RNA (piRNA) in yeasts (Honda et al., 2017). In elasmobranch genomes with abundant transposable elements, those tRNA loci may serve as components of the mechanism regulating the genomic readout. The expansion of intron-containing tRNAs in shark genomes (Fig. 3A), which modulate gene expression in yeasts (Nostramo et al., 2025), suggests a possible link to genome size variation. The enrichment of selenocysteine tRNA genes in shark genomes (Fig. 3A) underscores the need for future studies on selenium utilization in these species.

We also investigated a possible intragenomic bias of sequence length change and repetitive element distribution which potentially indicates selective factors. However, we did not detect a remarkable regional bias. First, we observed an increase in both intronic and intergenic sequence lengths along with chromosome lengths, while this trend is less pronounced in the red stingray (Fig. 4D, E). Second, elongated introns and intergenic sequences harbor a similar composition of repetitive elements to each other (Fig. 4G). This suggests that the maintenance of inserted repetitive elements during elasmobranch evolution did not have any marked regional preference.

Our study suggests that genome expansion is associated with increased gene duplication (Fig. 5D), possibly because larger genomes face reduced pressure to minimize the cost of additional genomic content, even if genes comprise only a small portion. Several previous studies on elasmobranch genomes examined gene family size changes, but they shed light on their association with particular biological processes including wound healing and longevity (Marra et al., 2019; Tan et al., 2021; Zhang et al., 2020; Yang et al., 2025). In our present study, we examined possible association of gene cluster size change with genome size, by employing a scoring method that is robust against the elongation of intergenic sequence length and inaccurate gene prediction (see Methods). Our genome-wide scan revealed larger tandem gene clusters in the red stingray with its larger genome, and this trend needs to be corroborated by more species with variable genome sizes, which is currently underway in the Squalomix consortium (Nishimura et al., 2022). The inventory of the identified tandem gene clusters included several gene families with increased gene copies in elasmobranchs, including SUCNR1 (Fig. 5B). SUCNR1 is a G protein-coupled receptor involved in the tricarboxylic acid (TCA) cycle, playing a crucial role in energy metabolism and physiological processes (He et al., 2004). Future research should explore how these duplications contribute to shaping the phenotypes of cartilaginous fishes.

Our analysis encompassed the genomic region containing the HoxC genes that was previously suggested to be absent in elasmobranch genomes (Jung et al., 2018; King et al., 2011). In the present study, we reported the retention of the red stingray HoxC genes, the first documentation among batoids which is evidenced by transcript identification (Fig. 6A; Supplemental Fig. S4B). The red stingray HoxC reduced its member but maintains a cluster, with high repeat frequency and elongated gene intervals (Fig. 6A), as demonstrated in orectolobiform shark genomes (Hara et al., 2018; Yamaguchi et al., 2023; reviewed in Kuraku, 2021; Kuraku, 2023). Shark HoxC clusters were identified on their X chromosomes, and its location was also maintained on the homologous red stingray chromosome, identified as ‘X1’, one of the two X chromosomes in this species (Niwa et al., 2025). On the other hand, our thorough search in the whitebelly skate genome and transcriptome identified no HoxC genes, confirming their absence in the entire order Rajiformes that was suggested consistently for other species within the order (Marlétaz et al., 2023). Our findings also indicate further genome reorganization in the myliobatiform lineage—in the red stingray genome, both HoxB and HoxC clusters are located on chromosome X1 (Fig. 6C). We speculate that the HoxC cluster underwent extensive repeat invasion following the adoption of the chromosome as a sex chromosome. Subsequent processes, possibly linked to genome size dynamics, may have led to the complete loss of the HoxC cluster in the rajiform lineage, which experienced marked genome compaction.

Our analysis revealed striking genomic differences between the red stingray and the whitebelly skate, including the scarcity of V2R and Pcdh gene cluster members and the complete absence of the HoxC gene cluster in the latter species. These differences, potentially influenced by genome size, could be linked to phenotypic differences, such as variations in electrolocation capacity (Last et al., 2016). Further comparative analyses, incorporating additional batoid lineages for which genomic data are still unavailable, will provide deeper insights into these patterns within broader taxonomic and ecological contexts.

## Methods

### Animals for genome sequencing

The female individual of the red stingray *Hemitrygon akajei* with the disc width of 37 cm (ToLID: sHemAka1) was captured in the Kinkai Bay, in Okayama, Japan. The female individual of whitebelly skate *Rhinoraja longicauda* with the disc width of 33 cm (ToLID: sRhiLon1) was captured off the coast of Sanriku, Japan (350m deep). Animals were handled, maintained, and the experiments were performed, in accordance with the Guidelines for Animal Experimentation established by Okayama University or Guideline of the Institutional Animal Care and Use Committee (IACUC) of RIKEN Kobe Branch (Approval ID: H16-11) in compliance with national regulations and international standards on animal welfare.

The remainder of the body of the individual was registered as a voucher specimen (ID: MNHAH A1111037) at the Museum of Nature and Human Activities, Hyogo (HITOHAKU), Japan for sHemAka1, and the residual body part of the whitebelly skate individual was registered as a voucher specimen (ID: SNFR 24105) in the collection of Japan Fisheries Research and Education Agency (FRA) (formerly Seikai National Fisheries Research Institute) for sRhiLon1.

### Genome sequencing and assembly

High molecular weight DNA was extracted from the liver of the red stingray and the heart of the whitebelly skate, using a NucleoBond AXG100 kit (Cat. No. 740545) of MACHEREY-NAGEL (Düren, Germany), which was followed by purification with phenol-chloroform. The concentration of the extracted DNA was measured with Qubit (ThermoFisher, MA, USA), and their size distribution was first analyzed with TapeStation 4200 (Agilent Technologies, CA USA) using the Genomic DNA ScreenTape (Agilent Technologies, CA USA) to ensure high integrity and later analyzed with pulse-field gel electrophoresis on CHEF DR-II (BioRad, CA, USA) to ensure the size range between 20 kb to 100 kb. To obtain long-read sequence data for each species, an SMRT sequence library was constructed with an SMRTbell Express Template Prep Kit 2.0 (PacBio, Menlo Park, CA, USA) and sequenced on a PacBio Sequel II system (PacBio). The sequencing output was processed to generate circular consensus sequences (CCS) to obtain a total of 149 Gb HiFi sequence reads in six SMRT cells for the red stingray and 106 Gb in four SMRT cells for the whitebelly skate. From these reads, adapter sequences were removed using the program HiFiAdapterFilt (Sim et al., 2022).

For the red stingray, the obtained HiFi sequence reads were assembled using the program hifiasm v0.16.1 (Cheng et al., 2021) with the option ‘-D 10’, resulting in contig sequences (sHemAka1.1). The obtained contigs were further scaffolded using optical mapping data obtained as follows. Genomic DNA was extracted from the spleen of the individual used for DNA sequencing with SP Tissue and Tumor DNA Isolation Kit (BioNano Genomics, Cat. No. #80038). DLE-1 was used for direct label staining. The labelling performed according to the BioNano Prep Direct Label and Stain Protocol. The labelled samples were scanned on the BioNano Saphyr system using Saphyr Chip G2.3 (BioNano Genomics, #20366), and the obtained data was processed with Access v1.7.1 (BioNano Genomics) followed by Solve v3.7_03302022_283 (BioNano Genomics) to perform pipeline analysis for *de novo* assembly followed by hybrid assembly. For the whitebelly skate, the obtained HiFi sequence reads were assembled using the program hifiasm v0.16.1 (Cheng et al., 2021) with the default parameters, resulting in contig sequences used in Hi-C scaffolding.

### Hi-C data production and genome scaffolding

Hi-C libraries for the red stingray and the whitebelly skate were prepared using the muscle and liver tissues of the individual used for DNA extraction, respectively, according to the iconHi-C protocol employing restriction enzymes DpnII and HinfI (Kadota et al., 2020). These Hi-C libraries were sequenced on a HiSeq X sequencing platform (Illumina Inc., CA, USA). The obtained Hi-C read pairs were processed with the program Trim Galore! v0.6.8 (https://www.bioinformatics.babraham.ac.uk/projects/trim_galore/) specifying the options ‘--phred33 --stringency 2 --quality 30 --length 25 --paired’ and aligned to the HiFi sequence contigs with the program Juicer v1.6 (Durand et al., 2016), and using its results, HiFi sequence contigs were scaffolded with 3d-dna version 201008 (Dudchenko et al., 2017) specifying the options ‘-m haploid -i 5000 --editor-repeat-coverage 15 -r 2’ to be consistent with the chromatin contact profiles.

The continuity of the genome assemblies, designated sHemAka1.3 and sRhiLon1.1, and completeness of the resultant gene models were assessed with the webserver gVolante v2.0.0 (Nishimura et al., 2017) in which the pipeline BUSCO v5.1.2 (Tegenfeldt et al., 2024) is implemented, consistently using the ortholog set ‘vertebrata_odb10’ supplied with BUSCO. The completeness of the genome assemblies was assessed with compleasm (Huang and Li, 2023). The genome assemblies were deposited in NCBI under the BioProject ID PRJNA1206076.

### Non-coding sequence annotation

The obtained chromosome-scale and other genomic sequences were subjected to the *de novo* detection of repetitive elements with RepeatModeler v2.0.4 (Flynn et al., 2020) with the -LTRStruct option. The detected repeat sequences were input in RepeatMasker v4.1.4 (Tempel, 2012) in the sensitive mode (with the option ‘-s’). Classification of the ten most abundant repeat elements in the red stingray genome was based on RepeatModeler. Simple tandem repeat detection was further reinforced by the use of the program tantan v40 (Frith, 2011). Putative transfer RNAs (tRNAs) were detected by the program tRNAscan-SE version 2.0.12 (Chan et al., 2021), and those with a cutoff score (‘-X’ parameter) of smaller than 50 were discarded. Detection of the rDNA-containing regions was performed with the program barrnap v0.9 (https://github.com/tseemann/barrnap) with an aid of the existing sequence entry spanning the oceanic whitetip shark rDNA locus (OP151210.1) and the human 45S pre-ribosomal sequence (NR_145819.1). Detection of canonical telomeric repeats (TTAGGG)n was performed with tidk v0.2.31 (Brown et al., 2023). For 5mC detection, subreads from which HiFi reads were derived for genome assembly was processed with ccsmeth (Ni et al., 2023) and the frequency of modifications was quantified in 1Mb non-overlapping windows, using the chromosome-scale genome assembly as a reference.

### Transcriptome sequencing and assembly

For transcriptome data acquisition, a young red stingray individual with a disc width of 15 cm was dissected to sample eye, liver, heart, stomach, midbrain, small intestine, gallbladder, pituitary, gill, ovary, kidney, and muscle. A red stingray embryo was also sampled, partly with dissection. For the whitebelly skate, the individual used for genomic DNA extraction was dissected to sample eye, liver, cerebellum, and telencephalon. These samples are listed in Supplemental Table S5. Total RNA extraction was performed using TRIzol reagent (ThermoFisher). After DNase I digestion, strand-specific RNA-seq libraries were prepared using 10-500 ng of each of the extracted total RNAs, with Illumina Stranded mRNA Prep kit (Illumina, Cat. No. 20040534) and IDT for Illumina RNA UD Indexes Set A Ligation (Illumina, Cat. No. 20040553) according to its standard protocol unless stated otherwise below. Before the total volume PCR amplification was performed, we performed a preliminary PCR using a 1.5 μL aliquot of 10 μL DNA from the previous step, with KAPA Real-Time Library Amplification Kit (Kapa Biosystems, Cat. No. KK2702). This demonstrated that the amplification of the products reached Standard 1 accompanying this kit between three and four PCR cycles, which instructed us to perform the full-volume PCR with three PCR cycles, introducing the minimal amplification. For embryonic transcriptome of red stingray, RNA-seq library was prepared based on Smart-seq v2 (Picelli et al., 2014). Input RNA was purified as described above and 1 ng total RNA was used for reverse transcription with a dT30VN primer (IDT). Oligo template switching was performed with mixing templated switching oligo (IDT), and reverse transcription was performed using SuperScript III (ThermoFisher, Cat. No. 18080044) with 10 cycles. cDNA was purified by adding 1.2x volume of AMpure beads, and cDNA was eluted with 10 μl of H2O. A 10 ng aliquot of cDNA was used for Tn5 tagmentation, and 5 cycles of PCR amplification were performed. The amplicon was purified by adding 1.2x volume of AMpure beads, and the final products were qualified using TapeStation 2200 with HS1000 detection kit (Agilent Technologies). The obtained sequence reads in the fastq files were processed with Trim Galore! v0.6.8 with the options ‘--phred33 --stringency 2 --quality 30 --length 25 --paired’. The reads after adaptor trimming were assembled with the program Trinity v2.15.1 with the options ‘--trimmomatic --SS_lib_type RF’.

### Gene prediction

Protein-coding genes were predicted with the program pipeline Braker3 (Gabriel et al., 2024) on the genome assembly sequences in which interspersed repeats and simple tandem repeats were soft-masked with a -nolow option with RepeatMasker v4.1.4 (see above) following RepeatModeler v2.0.4 as well as tantan v40 (Frith, 2011). The gene prediction incorporated all peptide sequences registered in the file ‘odb11_vertebrata_fasta’ provided by the OrthoDB database (Kuznetsov et al., 2023), as well as the output of paired-end RNA-seq read mapping with hisat2 v2.2.1 (Kim et al., 2019). In the genome assemblies of red stringray and whitebelly skate, this process yielded 19,725 and 16,649 predicted genes, respectively. Our assessment of the completeness of these gene sets using conserved one-to-one orthologs indicated the coverage of >95 %, resembling the completeness of the whole genome nucleotide sequences of >95 % measured by compleasm v0.2.6 (Huang and Li, 2023) (Supplemental Fig. S1).

### Molecular phylogenetic analysis

Protein sequences were collected from the NCBI and Ensembl databases, and their accession IDs used for the phylogenetic analysis are included in Supplemental Table S6. The deduced amino acid sequences were aligned with the MAFFT v7.505 (Katoh et al., 2019) using the L-INS-i method. The aligned sequences were trimmed with trimAl v1.4.rev15 (Capella-Gutiérrez et al., 2009) to remove unreliably aligned sites using the ‘-gappyout’ option. The maximum-likelihood tree was inferred with RAxML v8.2.12 (Hübner et al., 2021) using the PROTCATWAG model, and for evaluating the confidence of the nodes, the rapid bootstrap resampling with 100 replicates was performed. Molecular phylogenetic tree employing the Bayesian framework was inferred with PhyloBayes v4.1c (Rodrigue and Lartillot, 2014) using the CAT-WAG-Γ model.

### Detection of tandem gene duplications

Primary genome-wide survey of tandem gene clusters was performed by counting the number of adjacent genes judged as homologous by BLASTP v2.15.0+ (Altschul et al., 1997) with a threshold of E-value smaller than 1e-20 in ten neighboring protein-coding genes on both ends. For each predicted protein-coding gene, a number of neighboring homologous genes was scored between 0 and 20 (*M*_g_) (Fig. 5A; Supplemental Fig. S3). For fine-tune comparisons, the number and size of the tandem gene clusters detected in the primary survey were refined by manual inspection of nucleotide sequences of the genome assemblies.

### Hi-C data processing for centromere inference and TAD calling

For preprocessing, Hi-C data were processed using fastp v0.23.2 (Chen, 2023) to remove adaptor sequences and filter out low-quality reads with the options ‘-w 4 -f 1 -t 1 -l 30’. After preprocessing, the reads were mapped to the genome as single-end files using bowtie2 v2.5.4 with the options ‘-X 1000 --reorder --sensitive-local’. Mapped SAM files were converted to BAM format, indexed with default settings, and sorted by name (the option ‘-n’) using samtools v1.17 (Danecek et al., 2021). Hi-C matrix data was generated using HiCExplorer v3.7.5 (Ramírez et al., 2018; Wolff et al., 2020) with default settings. Restriction sites were identified using hicFindRestSite with the options ‘--searchPattern GATC’ for DpnII and ‘--searchPattern GANTC’ for HinfI. The Hi-C matrix was constructed using hicBuildMatrix v17 with the option ‘--minMappingQuality 15’. The output .h5 file was processed using hicCorrectMatrix with the option ‘--filterThreshold -1.5 5’.

Analysis of TADs and their boundaries were performed by binning the Hi-C matrices at resolutions of 10, 50, and 100 using hicMergeMatrixBins, which allowed the detection of TAD structures with varying resolutions. PCA analysis of the Hi-C data was performed inputting corrected .h5 files using hicPCA with the default setting. The outputs were visualized using hicPlotMatrix or pyGenomeTracks v3.8 (Lopez-Delisle et al., 2021; Ramírez et al., 2018).

## Data access

Raw sequence reads and the genome assembly have been deposited in NCBI under the BioProject ID PRJNA1206076. The individually annotated sequences are registered under accession IDs LC860076-LC860077.

## Acknowledgments

The authors thank Hidenori Nishihara, Kazuaki Yamaguchi, Osamu Nishimura, Kaori Tatsumi, Chiharu Tanegashima, Yuta Ohishi, Masayuki Iigo, Jérémy Berthelier, and Akiko Soma for insightful discussion, staff at Japan Fisheries Research and Education Agency for skate sampling, staff at Kazusa DNA Research Institute for long-read data acquisition, Susumu Hyodo for sharing samples, Hirofumi Harashima for optical mapping data acquisition, Tetsumi Takahashi at Museum of Nature and Human Activities, Hyogo, and Kouichi Hoshino at Fisheries Technology Institute, Japan Fisheries Research and Education Agency for assistance in voucher specimen registration. Computations were partially performed on the NIG supercomputer at the ROIS National Institute of Genetics.

## Author contributions

SK conceived the study. RM, TS, WG, WT, KSaito, provided materials. MK, SI, and KShirasawa acquired data. AK, YK, TN, SK performed analyses. All authors contributed to final writing of the manuscript.

