## Supplemental Info for "Tracing genome size dynamics in sharks and rays with inclusive sequence analysis by the Squalomix Consortium"

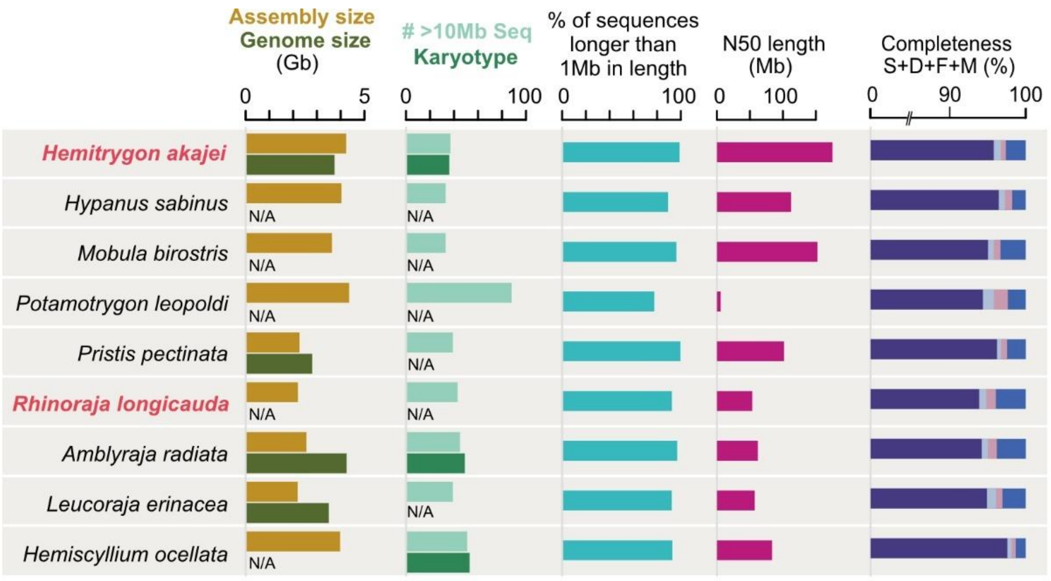


**Supplemental Figure S1.** Statistics of genome assembly in comparison with other species. See Methods for completeness assessment. Completeness scores are broken into the status of one-to-one ortholog identification: S (dark blue), single; D (light blue), duplicated; F (pink), fragmented, and M (navy), missing (see Methods).

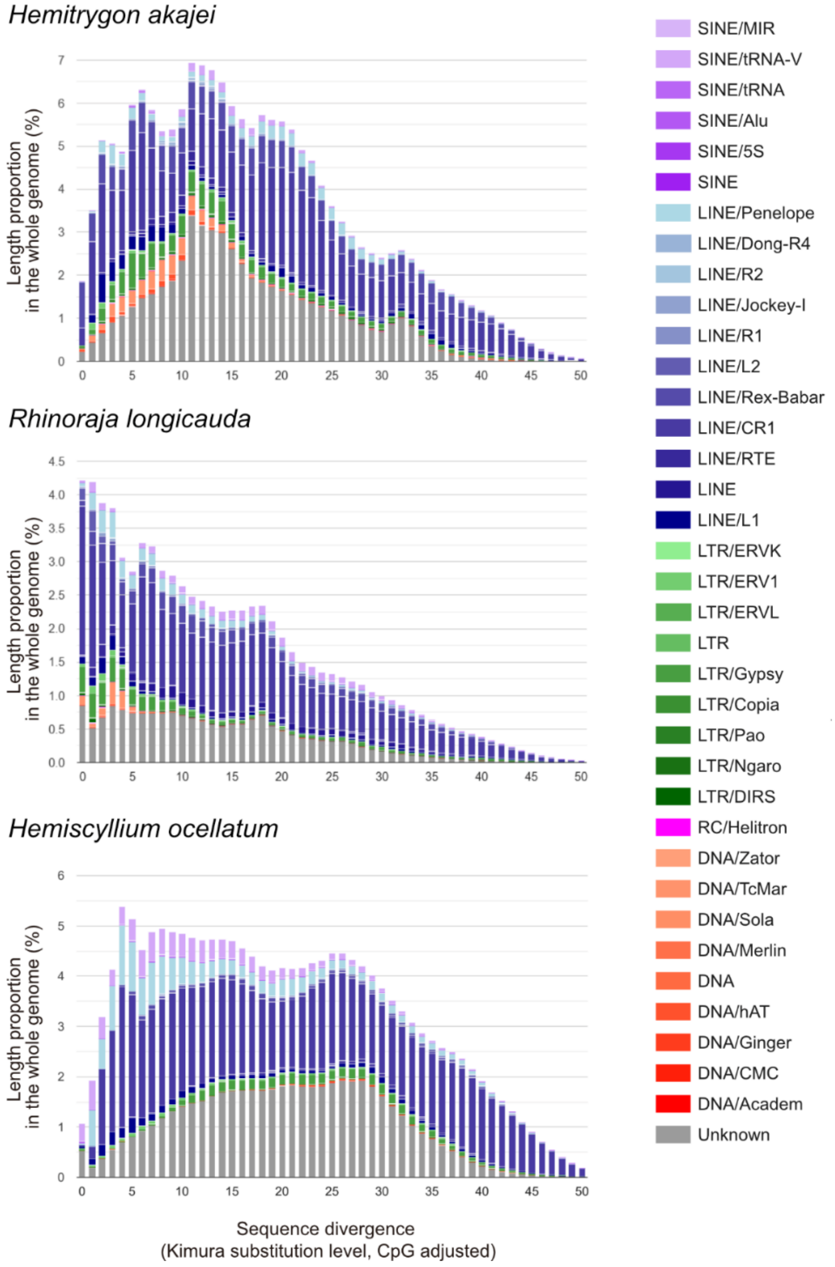


**Supplemental Figure S2.** Divergence distribution of interspersed repeats. The divergence of repetitive elements in individual repeat subclasses were computed using the Perl scripts, calcDivergenceFromAlign.pl and createRepeatLandscape.pl, that are accompanying the program RepeatMasker. Note that the vertical axes are not uniformly scaled. See Methods for more technical details.

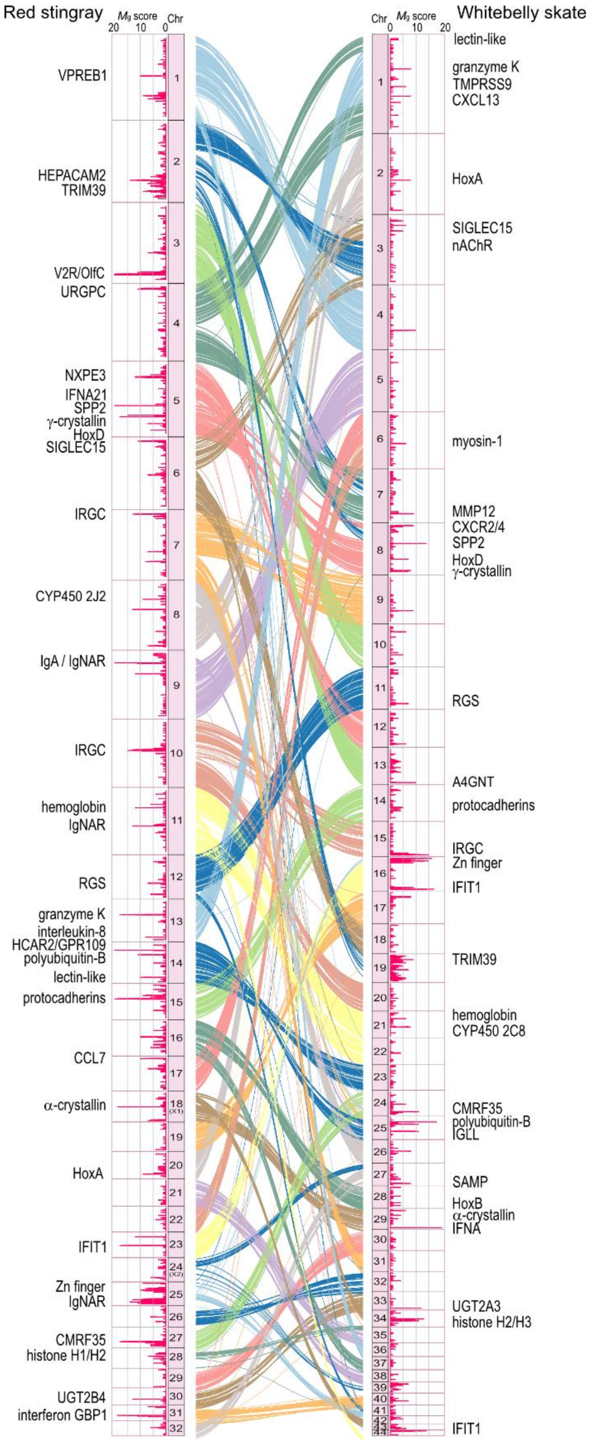


**Supplemental Figure S3.** Tandem gene clusters in batoid genomes. A, Landscape of tandem duplicates in the two batoid genomes produced in this study. The score $M_g$ (see Methods) for individual predicted genes is shown. The ribbons between the species show 1-to-1 orthology of protein-coding genes. Names of genes or gene groups are displayed when explicit non-elasmobranch homologs are identified. See Supplemental Table S4 for abbreviations of gene names.

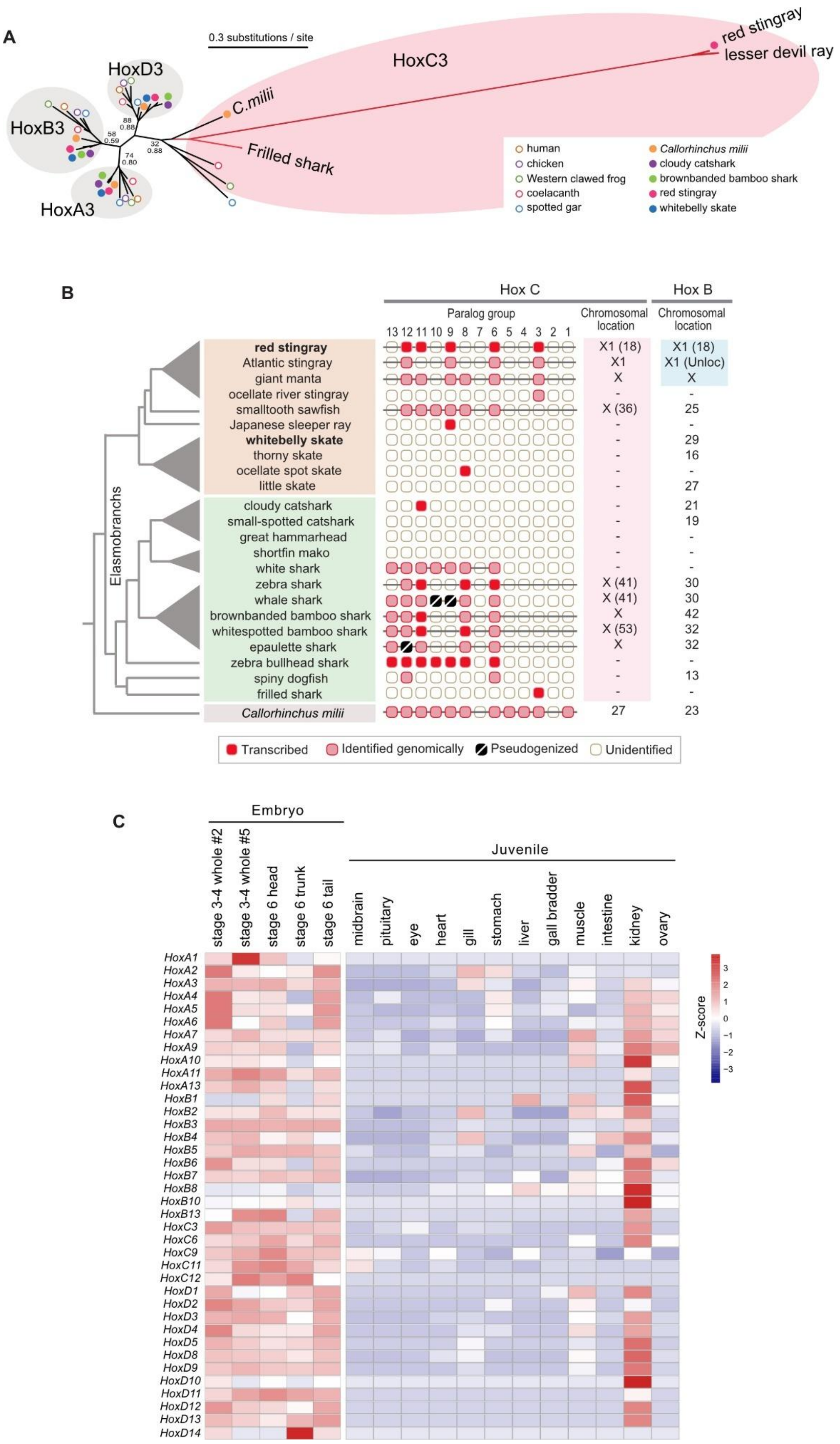
A
0.3 substitutions / site
HoxC3
red stingray
lesser devil ray
C.milii
Frilled shark
HoxD3
HoxB3
HoxA3
88
0.88
58
0.59
74
0.80
32
0.88
human
chicken
Western clawed frog
coelacanth
spotted gar
Callorhinchus milii
cloudy catshark
brownbanded bamboo shark
red stingray
whitebelly skate
B
Hox C
Hox B
Paralog group
Chromosomal location
13 12 11 10 9 8 7 6 5 4 3 2 1
Elasmobranchs
red stingray
Atlantic stingray
giant manta
ocellate river stingray
smalltooth sawfish
Japanese sleeper ray
whitebelly skate
thorny skate
ocellate spot skate
little skate
cloudy catshark
small-spotted catshark
great hammarhead
shortfin mako
white shark
zebra shark
whale shark
brownbanded bamboo shark
whitespotted bamboo shark
epaulette shark
zebra bullhead shark
spiny dogfish
frilled shark
Callorhinchus milii
X1 (18)
X1
X
-
X (36)
X (41)
X1 (Unloc)
X (53)
25
29
16
27
21
19
30
42
32
13
23
Transcribed
Identified genomically
Pseudogenized
Unidentified
C
Embryo
Juvenile
stage 3-4 whole #2
stage 3-4 whole #5
stage 6 head
stage 6 trunk
stage 6 tail
midbrain
pituitary
eye
heart
gill
stomach
liver
gall bradder
muscle
intestine
kidney
ovary
HoxA1
HoxA2
HoxA3
HoxA4
HoxA5
HoxA6
HoxA7
HoxA9
HoxA10
HoxA11
HoxA13
HoxB1
HoxB2
HoxB3
HoxB4
HoxB5
HoxB6
HoxB7
HoxB8
HoxB10
HoxB13
HoxC3
HoxC6
HoxC9
HoxC11
HoxC12
HoxD1
HoxD2
HoxD3
HoxD4
HoxD5
HoxD8
HoxD9
HoxD10
HoxD11
HoxD12
HoxD13
HoxD14
Z-score
3
2
1
0
−1
−2
−3

**Supplemental Figure S4.** Elasmobranch Hox B and C genes and their chromosomal locations. A, Detailed molecular phylogeny of Hox3 genes included in Fig. 6C. Red branches show the lineages leading to elasmobranch *HoxC3*. As support values for the ortholog groups, bootstrap probabilities in the ML tree are shown above posterior probabilities in Bayesian inference. See Supplemental. Table S6B for accession details for the sequences used. B, Repertoires of Hox C genes in chondrichthyans. Location of Hox B genes are also included. Gray triangles in the tree indicate order-level classification. Hyphens indicate the unidentification of the Hox cluster in genome sequences. C, Expression profiles of red stingray Hox genes based on RNA-seq. The heatmap shows $log_2$-transformed TPM values ($log_2$[TPM+1]) for selected genes across different tissues. The color gradient reflects the relative expression level, with blue indicating low expression and red indicating high expression. Expression values were scaled uniformly without row-wise normalization.

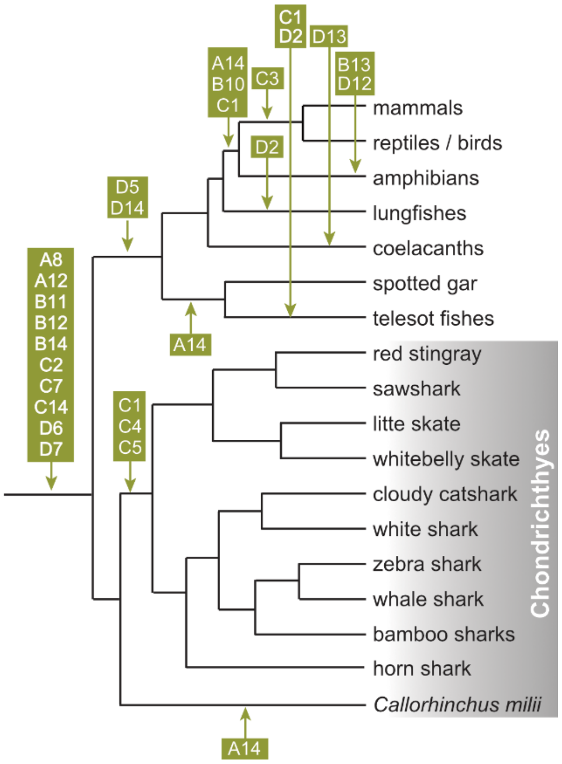


**Supplemental Figure S5.** Evolution of Hox gene repertoires. White letters indicate gene loss in individual evolutionary lineages. Loss of individual Hox genes, as well as the inference of their timings (arrows) are based on previous literature (Feiner and Wood, 2019; Kuraku and Meyer, 2009; Liang et al., 2011). Loss of HoxB and -C during the evolution of elasmobranchs are not indicated.

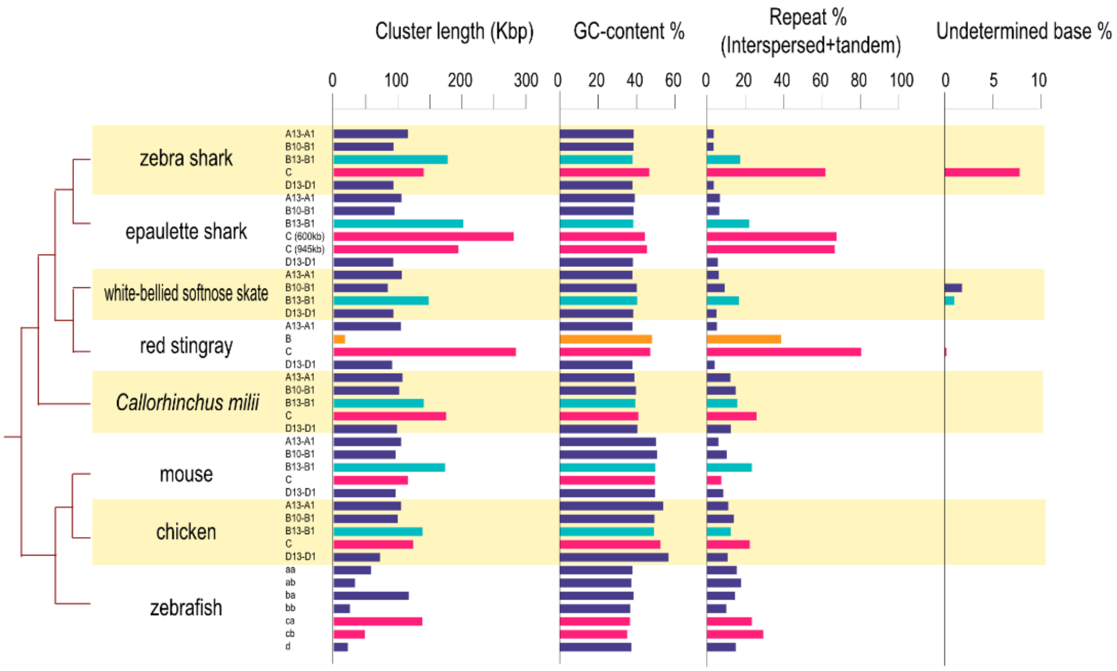


**Supplemental Figure S6.** Sequence properties of Hox clusters. The records presented here are converted to form Fig. 6C.

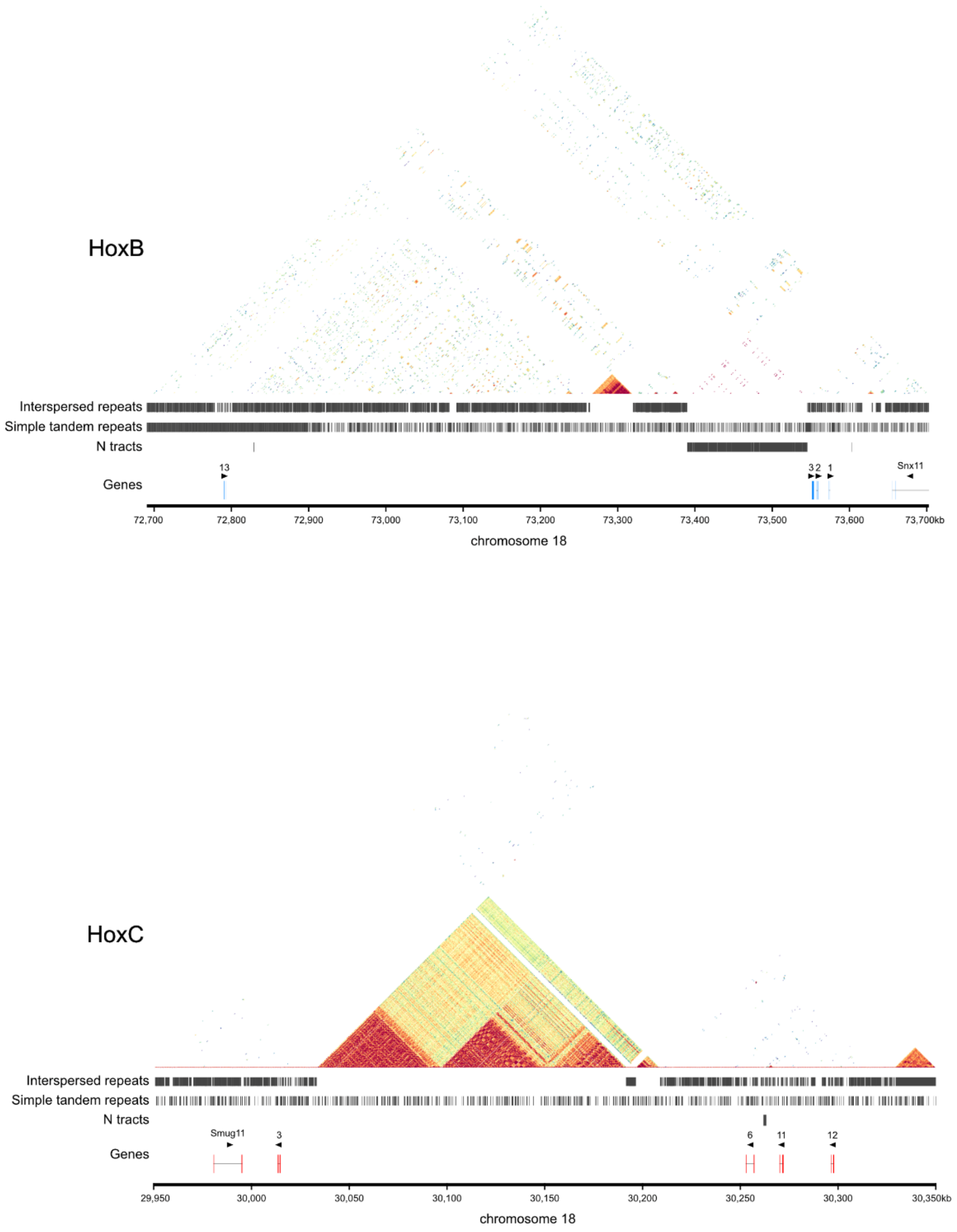


**Supplemental Figure S7.** Simple repeats distribution on the chromosomal sequences containing HoxB and -C. The HoxB-containing region harbors 17 undetermined regions including 15 consecutive gaps in the positions 73.4–73.55 Mb. The HoxC-containing region harbors one undetermined region.

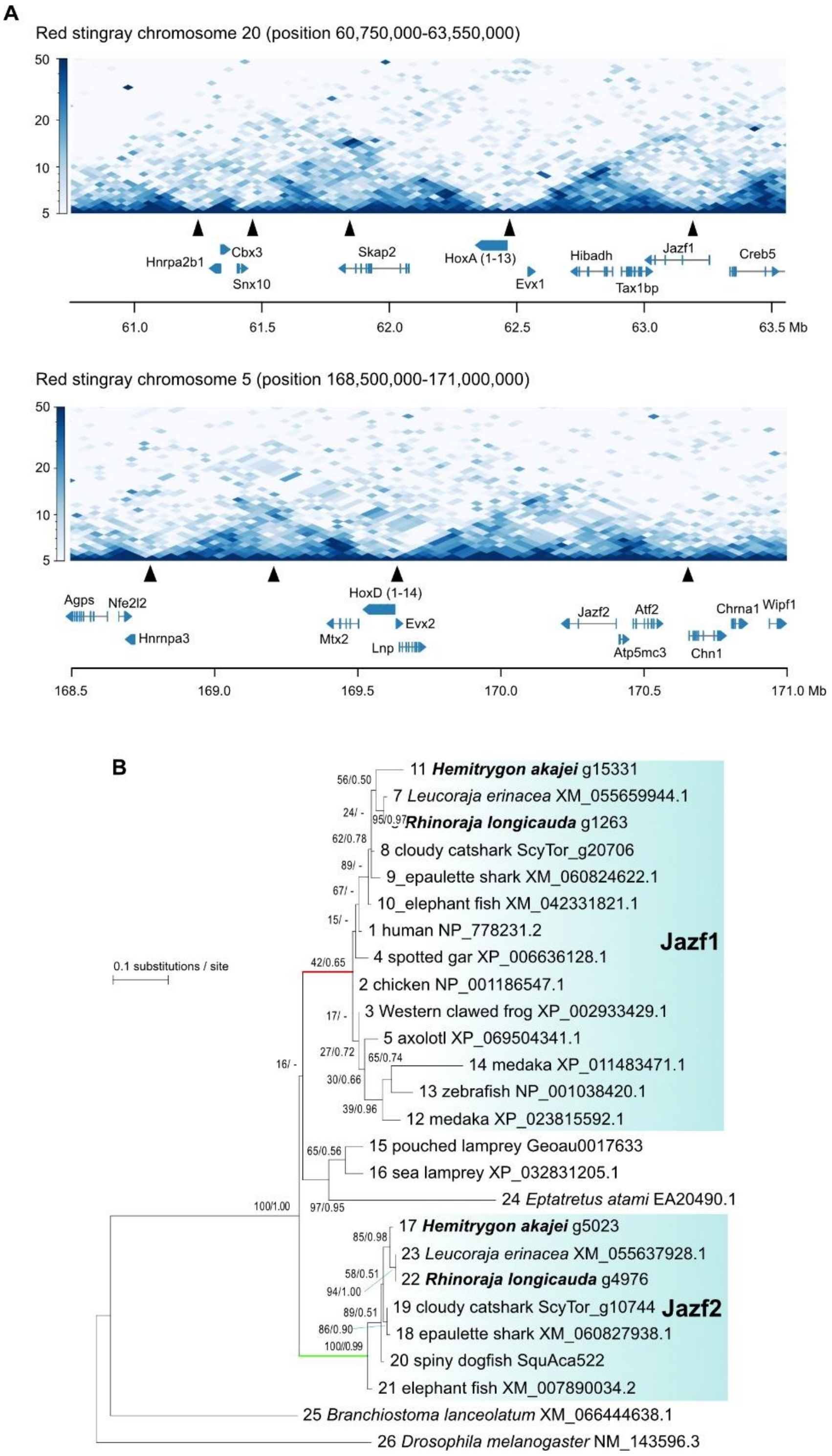


**Supplemental Figure S8.** HoxA and -D clusters and their surrounding regions. **A,** Topologically associating domains detected by Hi-C analysis. The Hi-C heat maps show chromatin contacts between different genomic regions near the Hox clusters. Arrowheads indicate the inferred boundaries of topologically associating domains (see Methods). The relative positioning of Hox clusters to two TADs—commonly referred to as C-DOM and T-DOM at HoxD cluster and as 3′ TAD and 5′ TAD at HoxA cluster—have been reported in other vertebrates, where the exact boundary location is influenced by biological contexts, such as tissue type and developmental stage (Andrey et al., 2013; Noordermeer et al., 2011; Rodríguez-Carballo et al., 2017; also see Results). **B,** Molecular phylogeny of Jazf1 (juxtaposed with another zinc finger 1) and Jazf2 genes shown in **A**. Jazf2, unrecognized previously in similar studies (Marlétaz et al., 2023), is a duplicate of Jazf1 gene and has been retained by cartilaginous fishes, while its osteichthyan ortholog is inferred to have been lost as no osteichthyan genes diverge from the branch shown in green in the tree. In contrast, Jazf1 is retained widely by different vertebrate lineages, while phylogenetic relationships within gnathostomes (inside the branch marked in red) are not inferred as expected from the widely accepted species phylogeny.

**Supplemental Table S1**. Systematic gap between the cytology-based and sequence-based measures of haploid genome size.

| Species | Common name | Experimentally measured nuclear DNA content (Gb) | Total length of genome DNA sequence assembly (Gb) |
|---|---|---|---|
| *Pristis pectinata* | smalltooth sawfish | 2.8 | 2.27 |
| *Amblyraja radiata* | thorny skate | 4.25 | 2.56 |
| *Leucoraja erinacea* | little skate | 3.5 | 2.19 |
| *Chiloscyllium punctatum* | brownbanded bamboo shark | 4.73 | 3.44 |
| *Chiloscyllium plagiosum* | white-spotted bamboo shark | 4.85 | 3.78 |
| *Scyliorhinus torazame* | cloudy catshark | 6.67 | 4.48 |
| *Scyliorhinus canicula* | small-spotted catshark | 5.34 | 4.22 |
| *Carcharodon carcharias* | white shark | 6.45 | 3.92 |
| *Squalus acanthias* | spiny dogfish | 5.8 | 3.71 |
| *Callorhinchus milii* | elephant fish | 1.9 | 0.94 |

Information in this table is visualized in the scatter plot in Figure 1B.

**Supplemental Table S2.** Interspersed repeats abundant in the red stingray genome.

| Identifier | Original length* (bp) | Length after curation (bp) | Repeat class** | GC-content (%) | Coverage in the whole genome*** (%) | Number of high-similarity copies**** |
|---|---|---|---|---|---|---|
| Erep16 | 4,100 | 4,150 | LINE/CR1 | 45.7 | 0.622 | 1,941 |
| Erep32 | 3,329 | 4,127 | LINE/CR1 | 49.7 | 0.037 | 3,146 |
| Erep68 | 2,668 | 4,208 | LINE/CR1 | 47.5 | 0.046 | 1,356 |
| Erep69 | 2,667 | 3,839 | LINE/CR1 | 47.1 | 0.152 | 1,811 |
| Erep92 | 2,221 | 2,461 | LINE/Penelope | 53.8 | 0.068 | 2,020 |

* Length of the sequence in the output of RepeatModeler.

**This classification is based on RepeatModeler.

***This count includes the genomic sequences with only short fractions of the elements detected by RepeatMasker.

****Genomic sequence stretches showing > 80% identity and >70% coverage of the query length. Those with <1000 copies are not included in this table.

**Supplemental Table S3**. Relationship between the intron length distribution and genome size.

| Species | Common name | Mean intron length (bp) | Genome size (Gb) |
|---|---|---|---|
| *Callorhinchus milii* | elephant fish | 2544 | 1.90 |
| *Rhinoraja longicauda* | whitebelly skate | 5216 | 2.19* |
| *Leucoraja erinacea* | little skate | 5401 | 3.38 |
| *Stegostoma tigrinum* | zebra shark | 7077 | 3.70 |
| *Squalus acanthias* | spiny dogfish | 7311 | 3.71* |
| *Hemitrygon akajei* | red stingray | 7353 | 3.74 |
| *Rhincodon typus* | whale shark | 6649 | 3.75 |
| *Hemiscyllium ocellata* | epaulette shark | 7579 | 3.98* |
| *Amblyraja radiata* | thorny skate | 5690 | 4.25 |
| *Chiloscyllium punctatum* | brownbanded bamboo shark | 7774 | 4.73 |
| *Scyliorhinus canicula* | small-spotted catshark | 11132 | 5.34 |
| *Scyliorhinus torazame* | cloudy catshark | 11608 | 6.67 |

Information in this table is visualized in the scatter plot in Figure 6A.

**Supplemental Table S4**. Gene families with tandem duplicates included in **Figure 6**.

| Abbreviation | Non-abbreviated name of gene groups |
|---|---|
| VPREB1 | V-set pre-B cell surrogate light chain 1 |
| HEPACAM2 | HEPACAM family member 2 |
| TRIM39 | E3 ubiquitin-protein ligase TRIM39 |
| V2R/OlfC | vomeronasal type 2 receptor |
| URGPC | upregulator of cell proliferation |
| NXPE3 | NXPE family member 3 |
| IFNA21 | interferon alpha-21 |
| SPP2 | secreted phosphoprotein 24 |
| SIGLEC15 | sialic acid binding Ig-like |
| IRGC | interferon-inducible GTPase 5 |
| CYP450 2J2 | cytochrome P450 2J2 |
| IgA/IgNAR | elasmobranch novel group (DELTA thalatoxin AVl1a-like) |
| RGS | regulator of G-protein signalling |
| HCAR/GPR109 | hydroxycarboxylic acid receptor/G protein-coupled receptor 109 |
| CCL7 | pC-C motif chemokine 7 |
| IFIT1 | interferon-induced protein with tetratricopeptide repeats 1 |
| CMRF35 | CD300C |
| UGT2B4 | UDP-glucuronosyltransferase 2B4 |
| interferon GBP1 | interferon guanine nucleotide-binding protein 1 |
| TMPRSS9 | transmembrane protease serine 9 |
| CXCL13 | C-X-C chemokine receptor type 13 |
| nAChR | nicotinic acetylcholine receptor |
| MMP12 | macrophage metalloelastase |
| CXCR2/4 | C-X-C chemokine receptor type 2/4 |
| A4GNT | alpha-1,4-N-acetylglucosaminyltransferase |
| CYP450 2C8 | cytochrome P450 2C8 |
| SAMP | serum amyloid P-component |
| UGT2A3 | UDP-glucuronosyltransferase 2A3 |

Note that gene groups names is based on the name of at least one of the human orthologs.

**Supplemental Table S5.** Properties of the sequencing libraries.

| SRA ID | Species | Data type | Library ID | Tissue |
|---|---|---|---|---|
| To be obtained | *Hemitrygon akajei* | HiFi reads | N/A | Liver |
| N/A | | BioNano Saphyr | N/A | Spleen |
| To be obtained | | Hi-C | P529_02_1 | Muscle |
| SRR32828577 | | RNA-seq | P341_01_1 | Right eye |
| SRR32828576 | | RNA-seq | P341_02_1 | Mid liver |
| SRR32828565 | | RNA-seq | P341_03_1 | Heart |
| SRR32828563 | | RNA-seq | P341_04_1 | Stomach |
| SRR32828562 | | RNA-seq | P341_05_1 | Midbrain |
| SRR32828561 | | RNA-seq | P341_06_1 | Small intestine |
| SRR32828560 | | RNA-seq | P341_07_1 | Gallbladder |
| SRR32828559 | | RNA-seq | P341_08_1 | Pituitary |
| SRR32828558 | | RNA-seq | P341_09_1 | Gill |
| SRR32828557 | | RNA-seq | P341_10_1 | Ovary |
| SRR32828575 | | RNA-seq | P341_11_1 | Kidney |
| SRR32828574 | | RNA-seq | P341_12_1 | Muscle |
| SRR32828573 | | RNA-seq | K0005_1 | Stage 6 embryo trunk |
| SRR32828572 | | RNA-seq | K0005_2 | Stage 3-4 embryo |
| SRR32828571 | | RNA-seq | K0005_3 | Stage 6 embryo tail |
| SRR32828570 | | RNA-seq | K0005_4 | Stage 6 embryo head |
| SRR32828569 | | RNA-seq | K0005_5 | Stage 3-4 embryo |
| To be obtained | *Rhinoraja longicauda* | HiFi reads | N/A | Heart |
| To be obtained | | Hi-C | P599_03_1 | Liver |
| SRR32828568 | | RNA-seq | P554_01_1 | Liver |
| SRR32828567 | | RNA-seq | P554_02_1 | Eye |
| SRR32828566 | | RNA-seq | P554_03_1 | Cerebellum |
| SRR32828564 | | RNA-seq | P554_04_1 | Telencephalon |

**Supplemental Table S6**. Sequences used for molecular phylogenetic tree inference.

A. SUCNR1

| Gene group | Species | Accession ID |
|---|---|---|
| Osteichthyan SUCNR1 | Human | NM_033050.6 |
| | Opossum | XM_056808671.1 |
| | Chicken | XP_025009233.1 |
| | Axolotl | XP_069476474.1 |
| | Western clawed frog | XP_017949524.1 |
| | Western clawed frog | XP_012818689.2 |
| | Western clawed frog | XP_017949523.1 |
| | Western clawed frog | XP_004914933.1 |
| | Western clawed frog | XP_031757720.1 |
| | medaka | XP_020563885.1 |
| | Spotted gar | XP_015216184.1 |
| Chondrichthyan SUCNR1cA | Little skate | XM_055645399.1 |
| | Little skate | XM_055646502.1 |
| | Red stingray | g2916.t1 |
| | Spiny dogfish | SquAca17896 |
| | Cloudy catshark | ScyTor_g5195 |
| | Cloudy catshark | ScyTor_g5196 |
| | Epaulette shark | XP_060690368.1 |
| | Epaulette shark | XP_060690010.1 |
| | Spiny dogfish | SquAca17897 |
| Chondrichthyan SUCNR1cC | Little skate | XM_055645750.1 |
| | Whitebelly skate | Manually curated* |
| | Red stingray | g2915.t1 |
| | Epaulette shark | XP_060690367.1 |
| | Cloudy catshark | ScyTor_g5197.t1 |
| | *Callorhinchus milii* | XM_007902499.2 |
| | Epaulette shark | XP_060690009.1 |
| | Cloudy catshark | ScyTor_g5198.t1 |
| Chondrichthyan SUCNR1cB | Little skate | XM_055645749.1 |
| | Red stingray | g2914.t1 |
| | Spiny dogfish | SquAca17898 |
| | *Callorhinchus milii* | XM_007902498.2 |
| | *Callorhinchus milii* | XM_007902495.2 |
| | *Callorhinchus milii* | XM_007902496.2 |

The order of the sequences in this table corresponds with that in Fig. 5B.

*This ORF was manually curated based on the nucleotide position 19809656-19808727 (reverse strand) on the scaffold 13.

B. Hox3

| Gene group | Species | Accession ID |
|---|---|---|
| HoxA3 | Spotted gar | XP_006636120.1 |
| | Human | NP_001371264.1 |
| | chicken | NP_001336689.1 |
| | Coelacanth | XP_005993141.1 |
| | Cloudy catshark | ASS31195.1 |
| | Brownbanded bamboo shark | g3556.t1 |
| | Red stringray | g15338.t1 |
| | Whitebelly skate | g1251.t1 |
| | *Callorhinchus milii* | XP_007898681.1 |
| | Western clawed frog | NP_001120901.1 |
| HoxB3 | Human | NP_001371676.1 |
| | Chicken | NP_001384824.1 |
| | Western clawed frog | NP_001015971.1 |
| | Coelacanth | XP_005990110.1 |
| | Spotter gar | XP_006638375.1 |
| | Cloudy catshark | ASS31206.1 |
| | Brownbanded bamboo shark | g7817.t1 |
| | Red stingray | DN67163_c7_g2_i1_2 |
| | Whitebelly skate | g13143+g13144* |
| | *Callorhnchus milii* | XP_007902552.1 |
| HoxC3 | Western clawed frog | XP_002936696.1 |
| | Coelacanth | XP_005995677.1 |
| | Spotted gar | XP_015199581.1 |
| | Red stingray | g14371.t1 |
| | Mobula | XP_062895325.1 |
| | Frilled shark | DN13016_c0_g2_i2** |
| | Callorhnchus milii | XP_007907853.2 |
| HoxD3 | human | NP_008829.3 |
| | chicken | XP_015145074.1 |
| | Western calwed frog | XP_002935734.1 |
| | coelacanth | XP_006000646.1 |
| | Spotted gar | XP_006636553.2 |
| | Cloudy catshark | ASS31216.1 |
| | Brownbanded bamboo shark | g5660.t1 |
| | Red stingray | g4999.t1 |

| | |
|---|---|
| Whitebelly skate | g4988.t1 |
| *Callorhnchus milii* | XP_007888242.2 |

*This ORF was retrieved by manually curating the sequences of the two separately predicted ORF with a support of transcript contigs.

**This ORF was identified in a transcript contig available at Squalomix BLAST Archive https://treethinkers.nig.ac.jp/squalomix/blast/ which is associated with our earlier study (Ohishi et al., 2023).
